# Alignment of LC-MS Profiles by Neighbor-wise Compound-specific Graphical Time Warping with Misalignment Detection

**DOI:** 10.1101/715334

**Authors:** Chiung-Ting Wu, David M. Herrington, Yizhi Wang, Timothy Ebbels, Ibrahim Karaman, Yue Wang, Guoqiang Yu

**Affiliations:** Department of Electrical and Computer Engineering, Virginia Polytechnic Institute and State University, Arlington, VA 22203, USA; Department of Internal Medicine, Wake Forest University, Winston-Salem, NC 27157, USA; Computational and Systems Medicine, Department of Surgery and Cancer, Imperial College London, London, SW7 2AZ, UK; Department of Epidemiology and Biostatistics, Imperial College London, London, W2 1PG, UK; UK Dementia Research Institute at Imperial College London, London, UK

## Abstract

**Motivation:** Liquid chromatography - mass spectrometry (LC-MS) is a standard method for proteomics and metabolomics analysis of biological samples. Unfortunately, it suffers from various changes in the retention times (RT) of the same compound in different samples, and these must be subsequently corrected (aligned) during data processing. Classic alignment methods such as in the popular XCMS package often assume a single time-warping function for each sample. Thus, the potentially varying RT drift for compounds with different masses in a sample is neglected in these methods. Moreover, the systematic change in RT drift across run order is often not considered by alignment algorithms. Therefore, these methods cannot completely correct misalignments. For a large-scale experiment involving many samples, the existence of misalignment becomes inevitable and concerning.

**Results:** Here we describe an integrated reference-free profile alignment method, neighbor-wise compound-specific Graphical Time Warping (ncGTW), that can detect misaligned features and align profiles by leveraging expected RT drift structures and compound-specific warping functions. Specifically, ncGTW uses individualized warping functions for different compounds and assigns constraint edges on warping functions of neighboring samples. Validated with both realistic synthetic data and internal quality control samples, ncGTW applied to two large-scale metabolomics LC-MS datasets identifies many misaligned features and successfully realigns them. These features would otherwise be discarded or uncorrected using existing methods. The ncGTW software tool is developed currently as a plug-in to the XCMS package.

**Availability and Implementation:** An R package of ncGTW is freely available at https://github.com/ChiungTingWu/ncGTW. A detailed user’s manual and a vignette are provided within the package.

**Supplementary information:** Supplementary data are available at Bioinformatics online.

## 1 Introduction

For proteomics or metabolomics analysis of biological samples, liquid chromatography coupled with mass spectrometry (LC-MS) is a standard method (Mueller, et al., 2007; Theodoridis, et al., 2008) that produces two-dimensional profiles of constituent compounds over retention time (RT) and mass-to-charge ratio (m/z). The identity and quantity of a particular compound (known (Lu, et al., 2008) or unknown (Vinaixa, et al., 2012)) may be inferred by analyzing the associated characteristic peak/curve profile (RT, m/z & intensity information). When analyzing multiple samples, the RT of each compound must be aligned accurately across different samples (Lu, et al., 2008).

Many software packages for LC-MS data analysis include a tool kit that performs RT alignment, including XCMS (Smith, et al., 2006), MZmine (Katajamaa and Oresic, 2005), and MS-DIAL (Tsugawa, et al., 2015). However, due to varying RT drift over different m/z bins in a sample and significant RT drift across distant samples (samples with larger run order difference), often nonlinear, accurate RT alignment remains a challenging task (Smith, et al., 2015). Unfortunately, classic alignment methods conveniently assume a single warping function across all m/z bins and perform multiple alignment that neglects the run order of each sample (Prakash, et al., 2006; Prince and Marcotte, 2006; Zhang, et al., 2005). These methods largely ignore the aforementioned two factors, and thus are prone to various types of misalignment. For a large-scale experiment involving many samples, the proportion of the misalignment becomes inevitable. For some types of misalignment, an algorithmic effort to mitigate them is to optimize parameter values in complex alignment algorithms. However, current strategies for handling a large number of parameters or features (peaks from the same compound at a single m/z bin with aligned RT across samples) are ad hoc, labor-intensive, subjective, and often fail to achieve a desired performance. Furthermore, for a misaligned feature whose true warping functions are different over different m/z bins, the misalignment cannot be corrected simply by adjusting the parameters of a single warping function (**Fig. 1**). Moreover, no existing analytics tool includes a systematic way to detect misalignment and thus previously acknowledged misalignment is often undetectable or uncorrected.

**Fig. 1.**
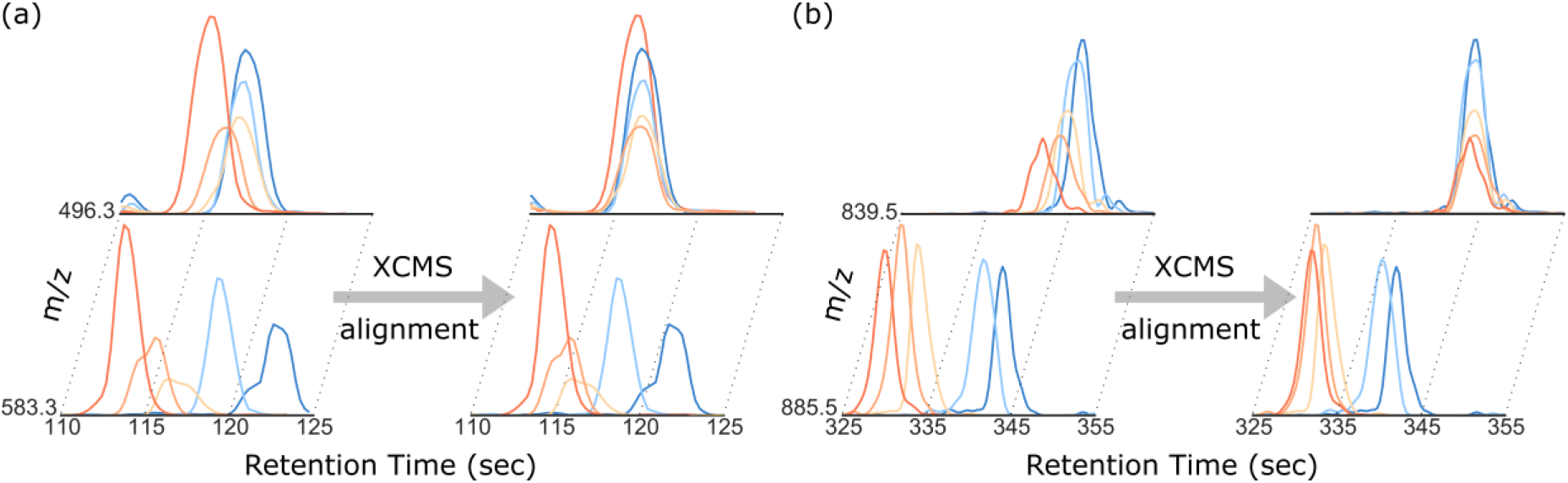
Examples of the observed misalignments due to single warping function assumption. Five samples over two m/z bins from each dataset are shown here for demonstration, where the upper and lower rows represent two different m/z bins respectively (see details in section 3.3). (a) An example from the Rotterdam dataset, shows that even with similar RT, the drift of each sample could be significantly different in two m/z bins. Using only a single warping function, XCMS can only align one bin (the upper one) well but not the other one as shown in the right part. (b) A similar example is also observed in MESA dataset.

To address the critical problem of the absence of validated methods for misalignment detection and structured alignment, we developed an integrated reference-free profile-based alignment method, neighbor-wise compound-specific Graphical Time Warping (ncGTW), that first detects misaligned features and then aligns affiliated profiles. In contrast to the feature-based methods that only align the detected peaks, fundamental to the success of our approach is the incorporation of expected RT drift structures across both different m/z bins or distant samples. Specifically, under the GTW framework (Wang, et al., 2016), ncGTW uses individualized warping functions for different m/z bins and assigns constraints on warping functions of neighboring samples. Furthermore, ncGTW utilizes a two-stage algorithm to achieve a reference-free alignment, where combinatorial pair-wise alignments are first performed and these aligned profiles are then coordinately aggregated into a pseudo-reference.

The input data to be analyzed by ncGTW includes the peak information extracted by XCMS and the raw data profiles corresponding to misaligned features. First, a statistically-principled misalignment detection scheme is applied to identify features requiring realignment. Then, each of the two-stage alignment procedures in ncGTW is solved efficiently by network flow based algorithms (Goldberg, et al., 2011). The ncGTW software tool takes full advantage of feature-based alignment methods and is provided currently as a plug-in to XCMS package.

## 2 Materials and methods

### 2.1 Detection of misaligned features

Accurate alignment of RT over a large number of samples remains a challenging task, particularly across distant samples due to significant yet varying RT drift. For a given feature corresponding to a set of peaks detected in several samples (e.g. by XCMS), we assume that the abundances of corresponding compound are independent of sample indices (this is an implication of randomizing the run order – standard practice in analytical science) and alignment relies on sufficient compound abundance in the relevant samples. Thus, for an accurately aligned feature, the samples associated with this feature should be a subset randomly drawn from the entire sample set, including both neighboring and very distant samples in the run-order domain (sample index). In contrast, a misaligned feature often splits into multiple features by the feature detection algorithm with non-ideal parameter setting (**Fig. 2**). In other words, when a detected feature includes only neighboring samples, it would highly likely constitute a misaligned feature (one of the most common forms of misalignment). Again, it is the very ability of being able to align distant samples merits an accurate alignment. Accordingly, we designed a statistically-principled approach to detect misaligned features. We associate the null hypothesis with correct alignment, use the range of sample indices within feature as the test statistic, and detect misaligned features by rejecting the null hypothesis.

**Fig. 2.**
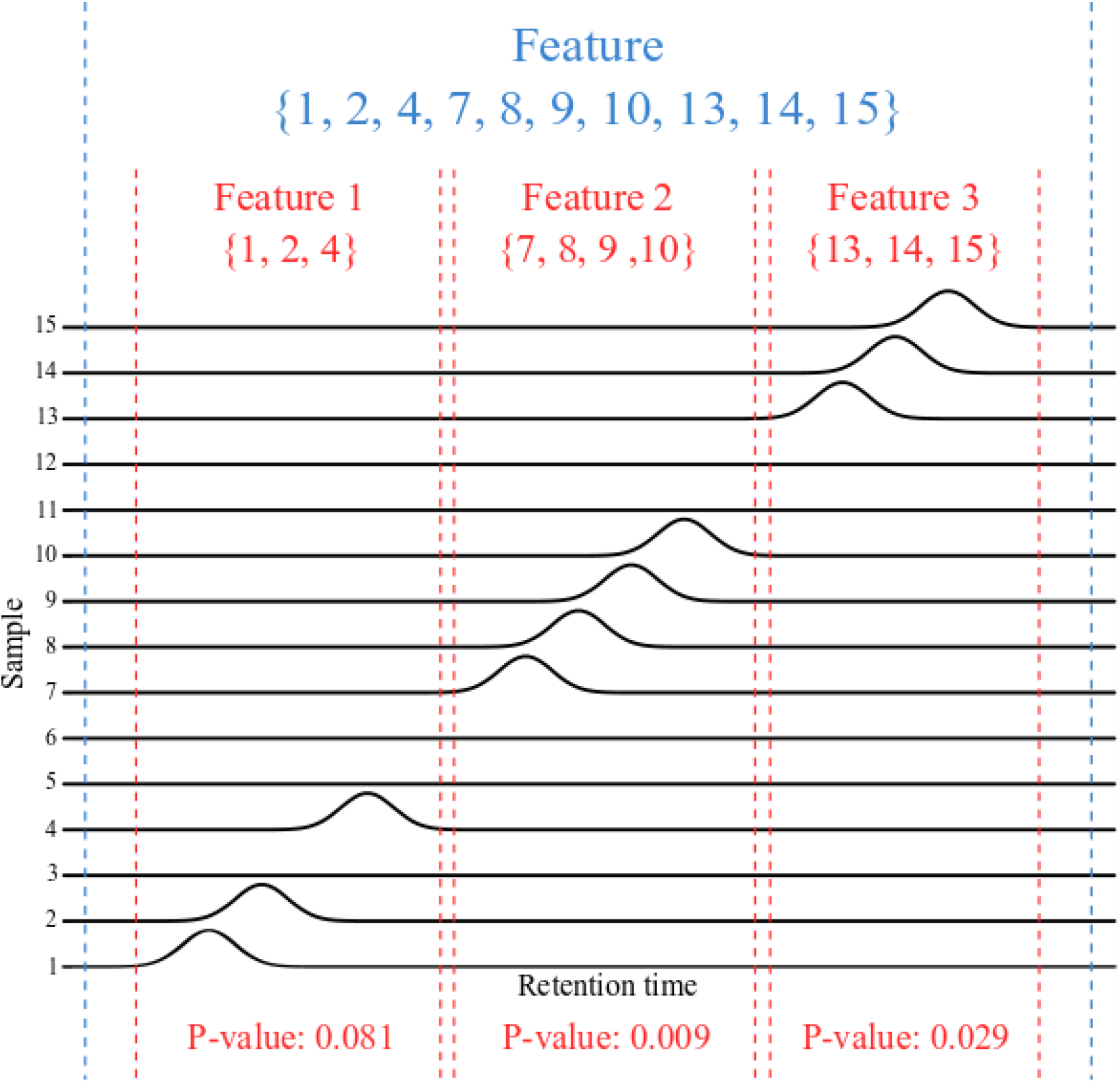
Illustrative example on detecting misaligned features. After initial alignment, among the total 15 samples, relevant peaks are detected only in samples 1, 2, 4, 7, 8, 9, 10, 13, 14, and 15, and some of the feature(s) are obviously misaligned. With a lower resolution grouping by XCMS, these peaks are all grouped into one single feature, as shown between the two blue dashed lines. While with higher resolution grouping, this feature is split into three features 1-3 as separated by the red dashed lines. The sample index sets of these features are {1, 2, 4}, {7, 8, 9, 10}, and {13, 14, 15} respectively. Only the p-values of features 2 and 3 are smaller than 0.05, thus pass the first criterion. Because the sample index sets of these two features are also disjoint, they pass the second criterion. Accordingly, ncGTW will realign the whole blue feature produced by the lower resolution grouping.

For a provisionally-aligned feature for which the peak is detected in a total of n samples, as the first criterion, we consider the range of sample indices in the feature as the test statistic, given by

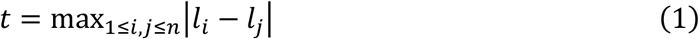

where *l*_*i*_ is the sample index of the *i*th sample in the feature, *n* − 1 ≤ *t* ≤ *N* − 1, and *N* is the total number of samples. Under the null hypothesis, we assume that *l*_*i*_ follows a discrete uniform distribution. Then, based on order statistics (Arnold, et al., 1992), it can be shown that the probability mass function of t is given by (Supplementary Information)

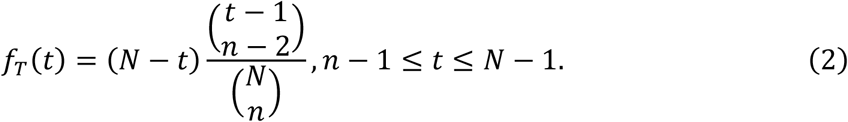

where *t* values associated with misalignment will always correspond to very small values under the null hypothesis, and the p-values associated with this feature can be estimated by

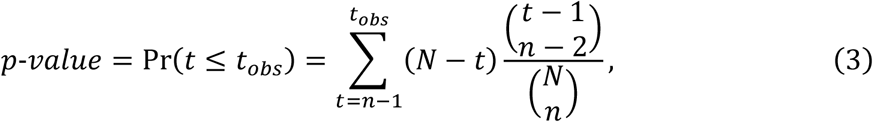

where *t*_*obs*_ is the observed sample index range of the feature. If the p-value of a feature is sufficiently small, reflecting the fact that the alignment only recruits neighboring samples, we can safely reject the null hypothesis and consider this feature as a candidate misaligned feature.

To address the additional layer of complication concerning varying RT drift over different m/z bins, for a candidate misaligned feature, we further check whether there exists neighboring feature(s) in the same m/z bin with sufficiently small p-value while with disjoint sets of sample indices, and if so, consider these features as needing to be realigned together. The rationale behind this second criterion is that applying same warping function cross different m/z bins would likely yet wrongly split the complete feature of same compound into several pieces.

In the ncGTW algorithm, these two criteria are combined to detect misaligned features. The initial alignment is performed using the existing XCMS alignment module. After XCMS alignment, two separate grouping results are produced using different RT window parameter values (bandwidth, bw) in XCMS peak-grouping module (feature detection). More precisely, the lower resolution grouping uses an RT window corresponding to the expected maximal RT drift, while higher resolution grouping uses RT window near to the RT sampling resolution (the inverse of scan frequency). First, ncGTW algorithm estimates the p-value of each feature using higher resolution grouping result and identifies all features with sufficiently small p-values and disjoint sample subsets. Then, the ncGTW algorithm matches the neighboring features to the corresponding features produced by lower resolution grouping, and considers realigning these features. An illustration of misalignment detection is given in **Fig. 2**.

The true causes for a misaligned feature may be complex and hidden, and may involve multiple yet unknown factors. It may be arguably suspected that the observed misalignment by existing alignment methods is at least partially due to the unstructured cost distributions over neighboring versus distant samples adopted by these methods. The joint optimization may thus be overly influenced by neighboring samples with intrinsically less costs. However, such biased solution is clearly in conflict with the very purpose of a globally optimal alignment that should be able to simultaneously correct larger and complex RT drift over distant samples.

### 2.2 Basic principles of profile-based multiple alignment

Different from peak-based alignment, profile-based alignment applies a gridded warping function to match two entire profiles with minimum cost. One popular method is pair-wise Dynamic Time Warping (DTW) (Sakoe, et al., 1990) that considers alignment as a shortest path problem. Specifically, DTW assigns cost to the edges of warping grids (**Fig. 3a**, the blue lines and dots) reflecting the dissimilarity between the two profiles being aligned. By the duality theorem, this shortest path problem can be readily converted into a graph of the maximum flow problem.

**Fig. 3.**
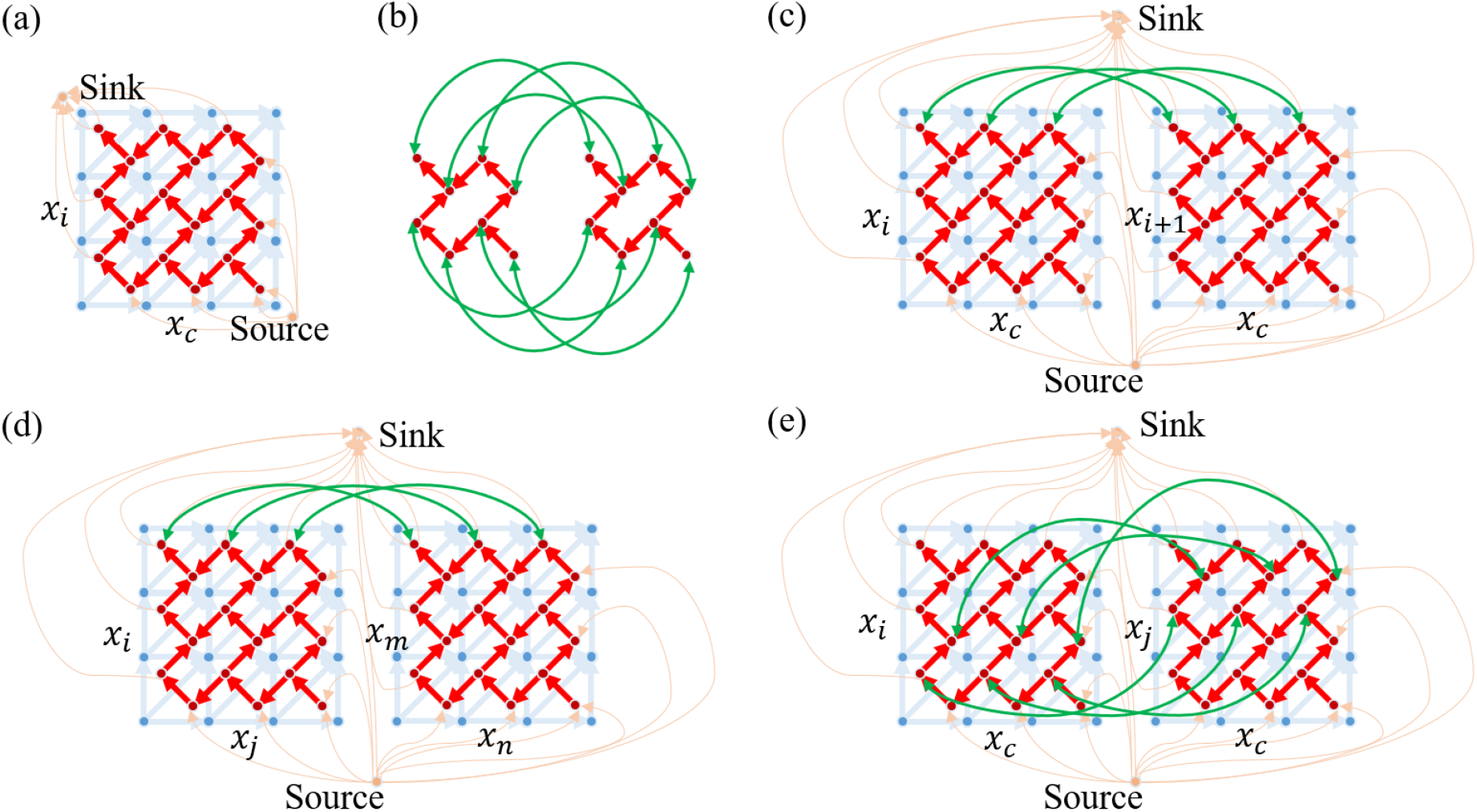
Construction of the various GTW graphs used in the different steps of ncGTW algorithm (Wang, et al., 2016). (a) DTW grid and corresponding graph for aligning a sample (*x*_*i*_) to a reference (*x*_*c*_), where the blue lines and dots form original DTW grid, and the red and orange lines and dots form corresponding maximum flow graph (DTW graph) (Wang, et al., 2016). Note here orange lines link only the vertices (those red dots enclosed by blue lined exterior triangle) to a single source or sink. (b) Two small DTW graphs are linked to form a GTW graph via additional ‘connecting’ edges (green lines between the corresponding vertices of two DTW graphs), where the weights on edges is a model parameter (Wang, et al., 2016). (c) GTW graph formed by two linked DTW graphs of two neighboring samples (*x*_*i*_ and *x*_*i*+1_) with a common reference, extendable to all neighboring samples, where the orange lines link vertices to a single source or sink forming a large maximum flow graph while green edges link the corresponding vertices of two DTW graphs (only edges linking top three vertices are shown here) (Wang, et al., 2016). (d) GTW graph constructed in Stage 1 of ncGTW without using a common reference, where *x*_*i*_ and *x*_*m*_ are neighboring samples, and *x*_*j*_ and *x*_*n*_ are neighboring samples. (e) GTW graph constructed in Stage 2 of ncGTW based on all pairwise warping functions obtained in Stage 1, where e.g., the warping function Φ_*i*→*j*_ guides the links between the corresponding vertices in GTW graphs, with *x*_*c*_ being the virtual reference.

To address some limitations of DTW, we have recently developed Graphical Time Warping (GTW) (Wang, et al., 2016) that extends classic DTW to performing multiple alignment with the flexibility of incorporating structure information such as neighboring versus distant samples in the run order domain (see more detailed discussions in (Wang, et al., 2016)). In graph theory, a flow network is defined as a directed graph involving a source and a sink and several other nodes connected with edges of limited capacity (Goldberg, et al., 2011). More specifically, after converting every DTW alignment grid to the graph of the maximum flow problem – DTW graph (**Fig. 3a**), additional edges are applied to link the DTW graphs (**Fig. 3b**) of neighboring samples and form a larger graph – GTW graph. These edges link the corresponding vertices of DTW graphs among neighboring samples (**Fig. 3c**), and the warping functions in the resulting larger graph are simultaneously estimated by solving the expanded maximum flow problem (Wang, et al., 2016).

In the application of GTW multiple alignment to real LC-MS data, we have also observed a few drawbacks of this method. Multiple alignment by GTW requires a common yet ‘ideal’ reference while in many real world problems, there is no a priori perfect reference available (e.g., no missing peak, little noise). Moreover, GTW may encounter a post-warping peak distortion problem.

### 2.3 Framework of ncGTW algorithm

To further address the drawbacks associated with GTW methodology, we developed the ncGTW framework that does not require a priori common reference and can correct potential post-alignment peak distortions. As mentioned above, GTW is quite sensitive to the selection of reference and performs well only if the reference choice is close to a good solution. Rather than requiring that a common reference be prespecified, the ncGTW algorithm takes a quite different approach and introduces a “circular multiple alignment” method that simultaneously considers all samples as references. The ncGTW algorithm consists of two key stages. First, by viewing each sample as a reference in a combinatorial network, ncGTW incorporates available sample structure (in the case of LC-MS data, the RT structure) information and aligns all possible sample pairs in the dataset without repetition. Then, ncGTW aligns all samples to a virtual reference using the warping point correspondences established in the first stage. The overall flowchart of ncGTW algorithm is given in **Fig. 4**.

**Fig. 4.**
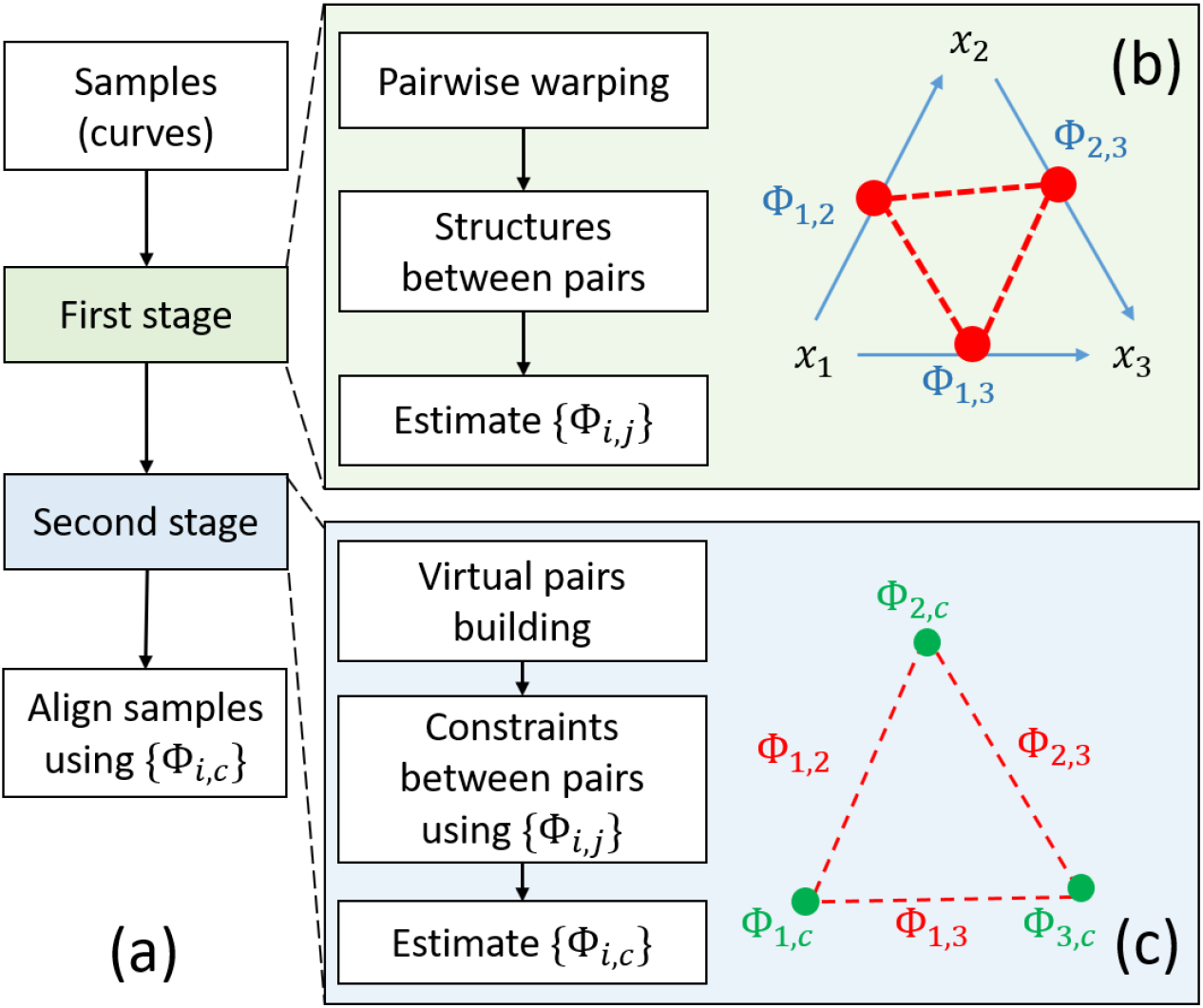
Flowchart of ncGTW algorithm. (a) With two-stage alignment strategy, all input samples (curves) are aligned simultaneously to a virtual reference. (b) Stage 1 of ncGTW with three illustrative samples. First, ncGTW builds a pairwise warping flow map (blue arrows). Then ncGTW incorporates structural information as the constraint and applies to all pairs (pair as red dot and constraint as a red dashed line). Lastly, ncGTW estimates all pairwise warping functions (Φ_*i*,*j*_) jointly with e.g., smoothness constraint on neighboring sample pairs. (c) Stage 2 of ncGTW with three illustrative samples. ncGTW aligns every sample to a common virtual reference *x*_*c*_, where the warping functions {Φ_*i*,*j*_} obtained in Stage 1 provide warping correspondences and final warping functions {Φ_*i*,*c*_} are calculated by solving the maximum flow problem.

In Stage 1, given the profiles {*x*_1_, …, *x*_*N*_} of *N* samples, ncGTW incorporates structural information and simultaneously estimates total *N*(*N* − 1)/2 warping functions {Φ_*i*,*j*_} for all distinct sample pairs. Like GTW, instructed by structural information, additional edges are applied to link the corresponding vertices of DTW graphs among neighboring samples (**Fig. 3d**). Again, this circular multiple alignment can be readily realized by solving classic maximal flow problem (Supplementary Information). Specifically, ncGTW estimates the enumerated pairwise warping functions jointly by minimizing the cost function given by

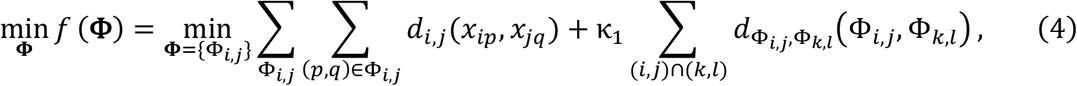

where 1 ≤ *i* < *j* ≤ *N*, (*p*, *q*) are the grid/point indices, *d*_*i*,*j*_ is the pointwise distance between sample *x*_*i*_ and sample *x*_*j*_, κ_1_ is a hyperparameter, (*i*, *j*) ∩ (*k*, *l*) indicates that the warping functions Φ_*i*,*j*_ and Φ_*k*,*l*_ are associated with neighboring samples, and 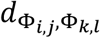 is the distance between each neighboring warping function pairs. All {Φ_*i*,*j*_} are efficiently obtained by solving an equivalent maximum flow problem.

In Stage 2, ncGTW uses the warping functions provided by {Φ_*i*,*j*_} as both guidelines and constraints to aggregate the profiles of all samples to a common virtual-reference, thus achieving reference-free multiple alignment. Note that the warping functions of ncGTW do not intentionally alter the point-wise intensity values of profiles but merely match the corresponding profile ‘coordinates’, that is, the cost of DTW graph edges in this stage has nothing to do with sample profile. In other words, the DTW graphs between samples and virtual-reference are ‘fully’ connected in the way that additional edges link all corresponding points of {Φ_*i*,*j*_} (**Fig. 3e**, green edges), where the cost of these connecting edges reflecting the distance between the corresponding warping points with respect to virtual reference. Furthermore, to prevent the possibility of all-to-one matching (all vertical/horizontal paths), a fixed cost is applied to only vertical/horizontal but not diagonal paths in the DTW graphs, encouraging warping functions with diagonal paths. Specifically, ncGTW estimates the final warping functions to virtual reference on all samples jointly by minimizing the cost function given by

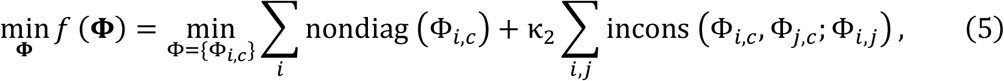

where nondiag (Φ_*i*,*c*_) is the cost measuring the deviation of warping function from diagonal path (one-to-one mapping), κ_2_ is a hyperparameter, and incons (Φ_*i*,*c*_, Φ_*j*,*c*_; Φ_*i*,*j*_) is the cost measuring the inconsistency between Φ_*i*,*c*_ and Φ_*j*,*c*_ given the pairwise warping function Φ_*i*,*j*_ (the corresponding points in Φ_*i*,*j*_ are aligned to different points on virtual reference *c*). Again, all {Φ_*i*,c_} are efficiently obtained by solving an equivalent maximum flow problem and thus the multiple alignment problem (Supplementary Information).

### 2.4 Correction of potential peak distortion

Peak distortion is a potential problem associated with DTW-based alignment, that is, while warping functions are optimally estimated by network flow algorithm, the shape of peaks may be altered. In other words, DTW-based methods do not guarantee one-to-one mapping for the points in the peak area (may be many-to-one or one-to-many), so the peak may become broader or narrower with shape changing. To avoid such distortion, based on multiple alignment of all samples to a virtual reference, within the starting and ending points of each peak, only the apex point will be aligned by the warping function and the rest of the points are simply shifted by the same amount as the apex. For the samples with missing peaks undetected by XCMS, the apices on virtual reference aggregated by other samples (no missing peaks) are first identified, then the corresponding points specified by warping functions are set as the apices of missing peaks in these samples, and lastly each ‘whole missing peak’ defined by the median of peak widths is aligned to virtual reference. An illustrative experimental result is given in **Fig. 5** using the MESA dataset, demonstrating the improved alignment by ncGTW where a misaligned feature with XCMS has been efficiently reported and corrected.

**Fig. 5.**
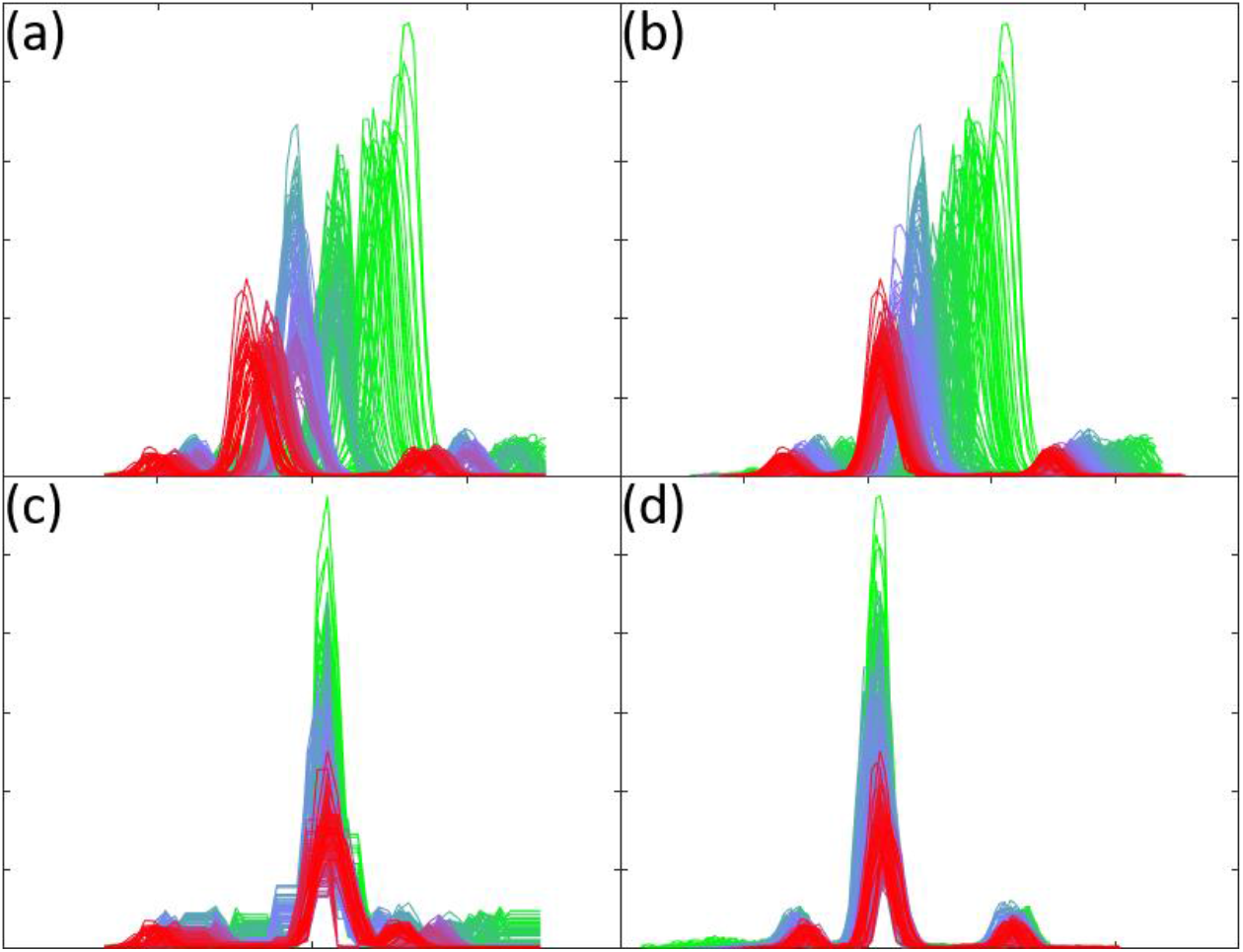
An illustrative experimental result on realignment and peak distortion correction by ncGTW, where a feature from the MESA dataset was initially misaligned by XCMS. The color mapping (green to blue and blue to red) corresponds to the sample index. (a) Raw LC-MS data associated with the feature of interest (before alignment). (b) The misaligned feature by XCMS that has been correctly detected and reported by the misalignment detection module of ncGTW package. (c) Realignment by ncGTW where apices are well aligned but with observable peak shape distortion. (d) Peak shape distortion is then efficiently corrected by the post-processing module of ncGTW package.

### 2.5 Integration of ncGTW into XCMS

The ncGTW R package is developed currently as a complementary plug-in to XCMS tool. The unique features of ncGTW algorithm include misalignment detection, individualized warping function (over m/z bins), incorporation of structural information (neighboring versus distant samples), reference-free multiple alignment, and correction of peak distortion. The functional integration of ncGTW into XCMS also allows ‘iteration’ (or ‘interaction’) between ncGTW and XCMS. For example, information about misaligned features obtained by ncGTW may be used to guide parameter retuning in XCMS. Then, those misaligned features may be realigned by ncGTW and regrouped by XCMS. Moreover, using the correct locations of missing peaks specified by ncGTW warping functions, those missing peaks may be accurately retrieved by the peak-filling procedure in XCMS. The workflow of XCMS-ncGTW pipeline is given in **Fig. 6**.

**Fig. 6.**
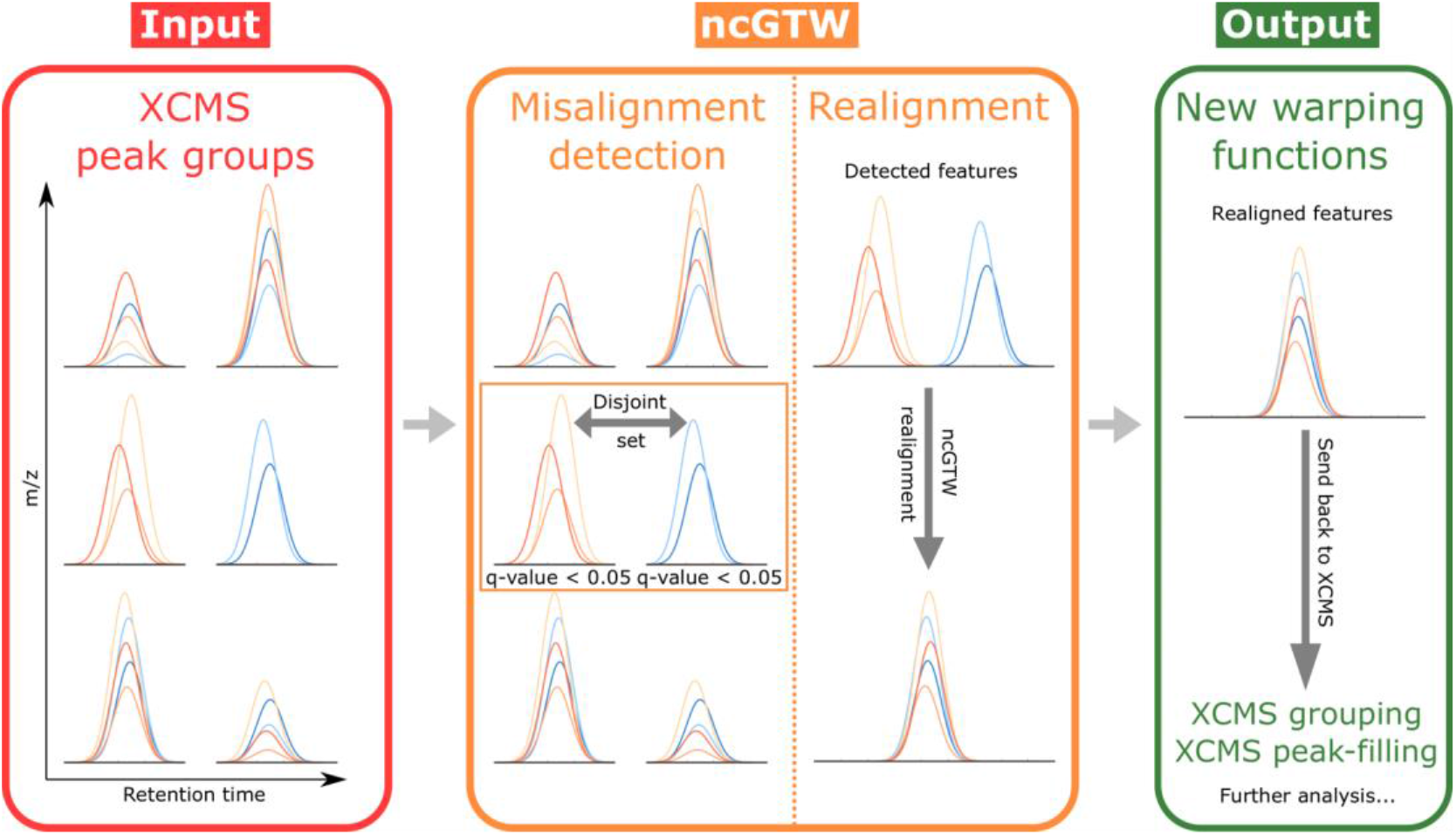
Workflow of ncGTW. As a plug-in to XCMS, ncGTW uses the grouping results provided by XCMS as the inputs (one lower resolution and one higher resolution, as explained in **Fig. 2**). Then, ncGTW detects all misaligned features using the aforementioned criteria and performs realignment on these features. Lastly, ncGTW calculates final warping functions for each sample that can be sent back to XCMS for re-grouping or peak-filling.

## 3 Results

### 3.1 Validation of ncGTW multiple alignment using simulated datasets

To test whether our ncGTW can improve multiple alignment by incorporating structural information of various forms, we assess alignment accuracy via simulation studies (Supplementary Information). Because the graph representation of structural information about neighboring samples is a unique feature of ncGTW, we consider three different structures including line (samples are profiled continuously), block (samples are profiled in batches), and uniform (no information on how samples are profiled) (Supplementary Information), in a set of experiments with 10 simulated samples. Three representative peer methods are selected for comparison including DBA (Petitjean, et al., 2011), CPM (Listgarten, et al., 2005), and GTW (Wang, et al., 2016). Specifically, DBA iteratively computes the barycenter of each aligned points set to give an average sample as a reference; CPM uses classic hidden Markov model to learn a prototype function for alignment, and GTW. Since GTW needs a reference, each sample is set as a reference and takes the average of scores in multiple alignment.

In addition to visual inspection, we use two benchmark quantitative measures to assess alignment accuracy, including mean correlation coefficient (MCC) and simplicity (SP) (Jiang, et al., 2013). MCC is averaged over all aligned sample pairs, and SP is the normalized sum of fourth power over all singular values on data matrix (Supplementary Information).

The experimental results are presented in Supplementary Information in detail and briefly summarized here. When there are missing peaks in some samples, DBA often misaligns some peaks, independent of specific neighborhood structure. When a selected reference contains missing peaks, GTW often produces many misalignments. CPM performs relatively better than DBA and GTW, while it misaligns several peak groups with block structure of large drift. The proposed ncGTW approach consistently outperforms all three peer methods by aligning all the peaks with the highest scores in all accuracy measures and neighborhood structures. **Table 1** gives the average performance scores obtained in simulation studies.

**Table 1.** The average performance scores obtained in the simulation studies with line, block, and uniform structure, where ‘Before alignment’ serves as the baseline, MCC and SP represent mean correlation coefficient and simplicity respectively, and the range of either score is between 0 and 1 with higher score indicating better performance.

| **Line structure** |  |  |
| --- | --- | --- |
| Scores<br>Methods | MCC | SP |
| Before alignment | 0.2330 | 0.4202 |
| DBA | 0.5422 | 0.6737 |
| CPM | 0.7894 | 0.8981 |
| GTW | 0.7991 | 0.9592 |
| ncGTW | **0.8366** | **0.9999** |
| **Block structure** |  |  |
| Scores<br>Methods | MCC | SP |
| Before alignment | 0.1062 | 0.1959 |
| DBA | 0.7767 | 0.8761 |
| CPM | 0.3773 | 0.5078 |
| GTW | 0.8687 | 0.9521 |
| ncGTW | **0.9159** | **0.9998** |
| **Uniform structure** |  |  |
| Scores<br>Methods | MCC | SP |
| Before alignment | 0.0641 | 0.2341 |
| DBA | 0.3797 | 0.7649 |
| CPM | 0.5651 | 0.8867 |
| GTW | 0.4926 | 0.8447 |
| ncGTW | **0.6953** | **0.9990** |

### 3.2 Initial test of ncGTW multiple alignment on small-scale real LC-MS dataset

We further conduct similar comparison studies on a small-scale real LC-MS dataset of 10 samples involving line, block, and uniform structures. Because no ground truth on correct alignment is available, we opt for visual inspection to assess relative performance. The experimental results are highly consistent with what is observed in simulation studies. DBA and GTW easily misaligned samples with some missing peaks, and CPM failed again on either block structure or samples with significant peak intensity imbalance. In contrast, ncGTW aligned all peaks nicely and consistently under different neighborhood structures. The alignment results are shown in **Fig. 7** (with line structure), and additional experimental results using other neighboring structures are given in Supplementary Information.

**Fig. 7.**
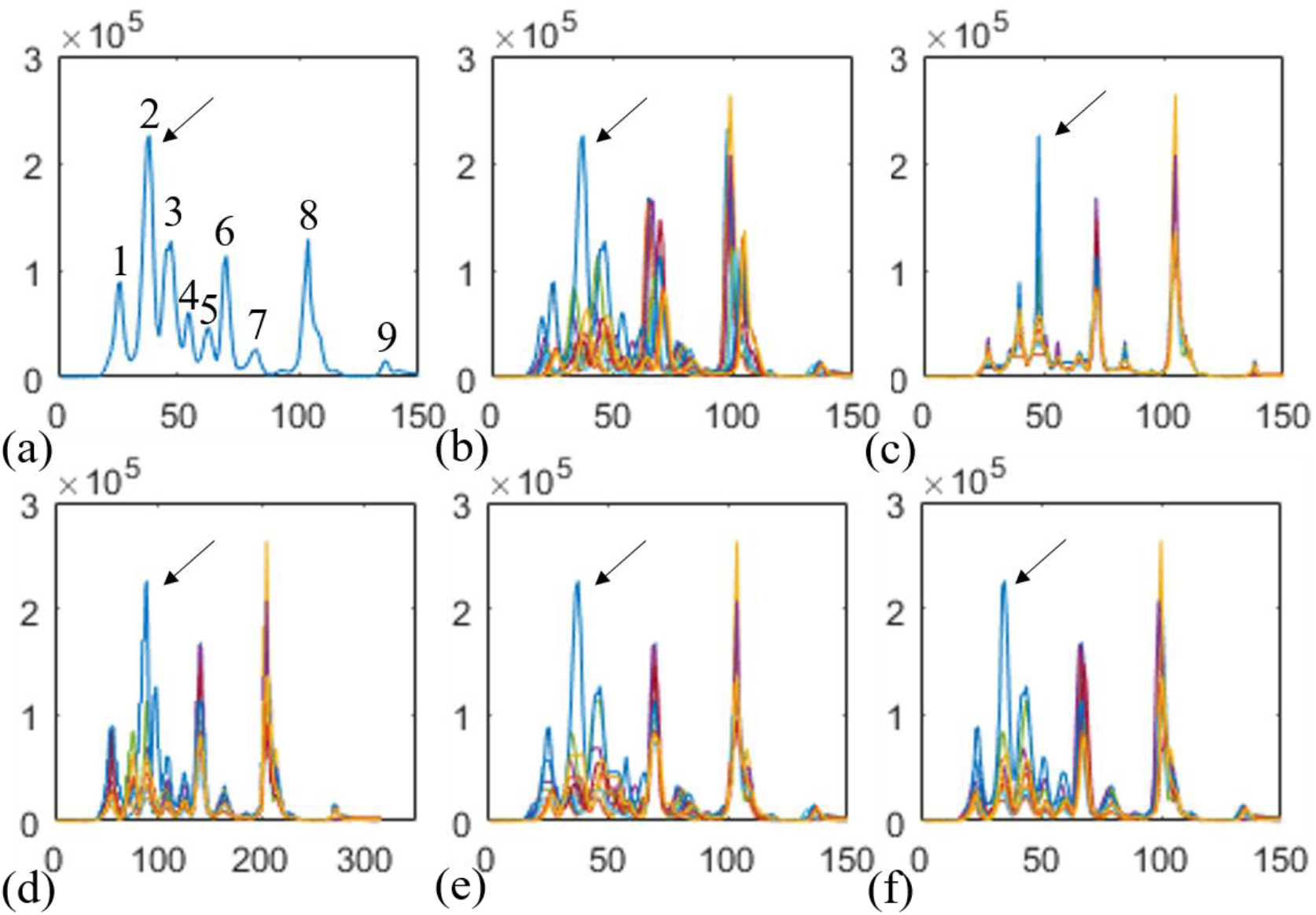
Illustrative realignment successfully performed by ncGTW incorporating “line” structure in small-scale real LC-MS dataset. (a) LC-MS profile with nine indexed peaks where the second peak (indicated by arrow) was misaligned by most peer methods except GTW and ncGTW. (b) The same LC-MS profiles of total ten samples. (c) The arrow-indicated second peak was wrongly aligned to the third peak-group (and all peaks were severely distorted by DBA). (d) The arrow-indicated peak was misaligned by CPM. (e) The arrow-indicated peak was well-aligned by GTW while the fourth and fifth peak-groups were misaligned. (f) All nine peak-groups were correctly and accurately aligned by ncGTW in this challenging case.

### 3.3 Description of Rotterdam and MESA cohorts and experimental settings

We apply ncGTW pipeline to two large-scale real metabolomics LC-MS datasets, namely the Rotterdam (The Rotterdam Study, The Netherlands) and MESA (The Multi-Ethnic Study of Atherosclerosis, USA) cohorts (Bild, et al., 2002; Hofman, et al., 2013), first to detect misalignment and then to realign those misaligned features.

Experimental protocols for reversed-phase liquid chromatography used by the National Phenome Centre (UK) were followed for data acquisition (Lewis, et al., 2016). LC-MS profiles of the serum samples from the Rotterdam and MESA cohorts were generated using a Waters Acquity Ultra Performance LC system (Waters, Milford, MA, USA) for chromatographic separation and a Xevo G2S Q-TOF (Waters, Milford, MA, USA) for m/z separation in positive ionization mode.

The Rotterdam dataset contains 1,000 study samples and 44 internal quality control (iQC) samples and the MESA dataset contains 1,977 study samples and 335 iQC samples. Each iQC sample is an aliquot of a pool of all the study samples, used to monitor and correct instrument performance in long runs. Because each of these two cohorts contains many samples (> 1k), the total time duration on data acquisition would be in the range of weeks - thus significant RT drift across experiments is expected. Indeed, on the iQC samples, the global warping function assumption made by XCMS is clearly and evidently violated in both Rotterdam (**Fig. 1a**) and MESA (**Fig. 1b**) datasets (using five samples from each dataset for demonstration). Moreover, in our observation, we estimate that there are around 3% of features which are misaligned due to the global warping function assumption in each dataset. Given that these two datasets were acquired from different cohorts at different labs with probably different experimental settings, this kind of misalignment may happen in any large dataset, and is not a rare occurrence.

In the subsequent experiments on both iQC and study samples, the major preprocessing steps (peak detection, RT alignment, peak grouping) are done by XCMS, and the ncGTW pipeline uses the default parameter settings.

### 3.4 Detection of misaligned features in Rotterdam and MESA datasets

Application of XCMS to the Rotterdam iQC samples alone generates total 1,872 features, among which 57 features are detected as potentially misaligned by the misalignment detection module in ncGTW package. With a closer visual inspection (performed independently by two MS experts), 41 features are confirmed as misaligned (true positives, **Fig. 5b** as an example), and 16 remaining features are considered well-aligned (false positives, see **Fig. S11** as an example). On Rotterdam study samples (excluding iQC samples), XCMS generates total 1,689 features, of which 45 features are detected as misaligned. Visual screening identifies 32 true positives and 13 false positives. On MESA iQC samples alone, XCMS generates total 1,951 features, of which 61 features are detected as misaligned. Visual inspection identifies 58 true positives and 3 false positives. On MESA study samples (excluding iQC samples), XCMS generates total 1,861 features, of which 49 features are detected as misaligned. Visual screening identifies 48 true positives and 1 false positive. While the false discovery rate is higher than theoretically expected, these false positives are mainly due to signal intensity fluctuation and will be eliminated in a post-realignment step.

### 3.5 Realignment by ncGTW

We apply the ncGTW tool to realign the features flagged as misaligned in the previous step. While we have applied ncGTW to all study samples to show the scalability, our reports are focused on the results only on iQC samples mainly attributed to the feasibility of performing quantitative assessment. We use two benchmark quantitative measures to assess the comparative performances by ncGTW and XCMS. The evaluation criteria are the average pairwise correlation coefficient (Jiang, et al., 2013) and the average pairwise total overlapping area (Christin, et al., 2008).

Comparative experimental results are detailed using two performance measures (**Fig. 8**), where the features with circles represent true positives and the features with crosses represent the false positives. These comparative experimental results, on both Rotterdam and MESA datasets, consistently show that ncGTW effectively and accurately realigns those misaligned features. Specifically, ncGTW realignment on most true positives achieves much higher performance scores than that of XCMS, and on most false positives, produces comparable performance scores as expected (near the diagonal lines, **Fig. 8**). Note that the default parameter setting of ncGTW in this step is purposely designed for the true positives, and probably not suitable for the false positives.

**Fig. 8.**
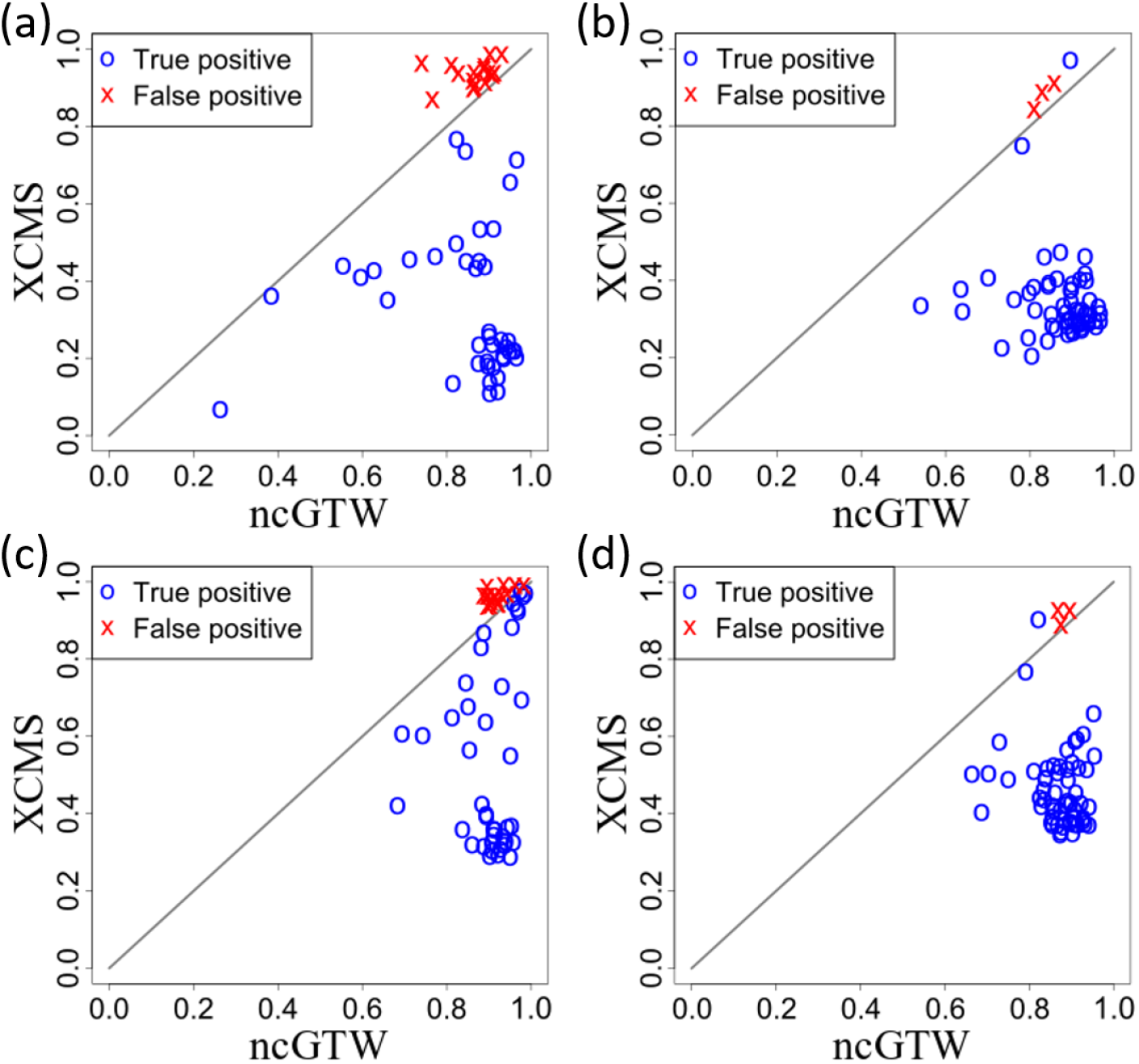
Application of ncGTW realignment method to Rotterdam and MESA datasets, where among the detected misaligned features, the blue circles represent true positives, and the red crosses represent false positives, respectively. (a) The average pairwise correlation coefficients on the Rotterdam dataset. (b) The average pairwise correlation coefficients on the MESA dataset. (c) The average pairwise total overlapping area on the Rotterdam dataset. (d) The average pairwise overlapping area on the MESA dataset.

Experimental results on iQC samples also show that the two performance scores can well separate the true positives and false positives (**Fig. 8**). Accordingly, via a post-alignment step we use these two performance scores to screen out true positives for further analysis. This strategy is also applicable to handling study samples.

### 3.6 Evaluation of ncGTW via post-realignment peak-filling performance

Accurate alignment of RT drift has significant impact on the performance of peak-grouping and peak-filling that define the features. In the XCMS pipeline, detected peaks are first grouped into features, and when there are undetected/missing peaks, peak-filling is then performed to retrieve those peaks. In our experiments, the coefficient of variation (CV) of intensity calculated over the samples, with and without ncGTW guided peak-filling, is adopted to assess the beneficial impact of ncGTW realignment (Matuszewski, et al., 1998).

Though we proposed that we can screen out the false positives by the two performance scores, we still include the false positives in the peak-filling step to observe the impact of the realignment on them. Post-realignment peak-filling results show that, measured by CV over the iQC sample features, ncGTW realignment consistently reduces the CV as compared with that derived from the initial XCMS alignment in both Rotterdam and MESA datasets (**Fig. 9**). Note that here again the circles represent the true positives and the crosses represent the false positives. More importantly, post-realignment peak-filling supported by ncGTW has led to the significantly reduction of CV on many ‘hard-to-define’ features, demonstrating the beneficial contribution of ncGTW realignment to improved feature generation.

**Fig. 9.**
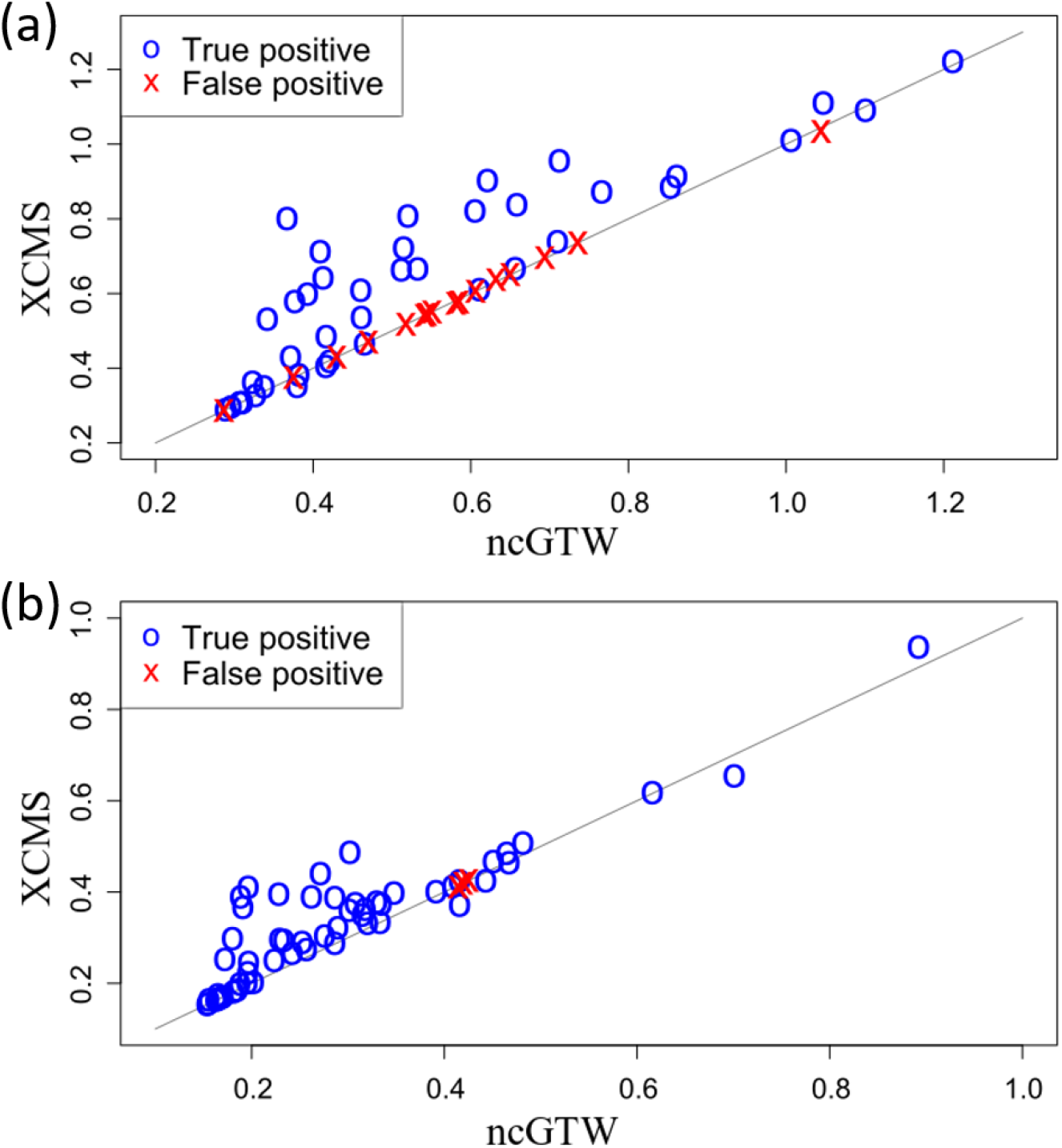
The comparisons of CV with versus without ncGTW realignment after the peak-filling step of XCMS. The blue circles represent the true positives, and the red crosses represents the false positives. (a) The CV comparison on Rotterdam dataset. (b) The CV comparison on MESA dataset.

## 4 Discussion

Feature-based and profile-based alignment methods are complementary to each other. Because of high efficiency on large datasets, the majority of existing alignment methods are feature-based. However, due to the challenging nature of accurate peak detection particularly when there are some missing peaks and significant RT drift, misalignment occurs on some features that also causes incorrect peak-grouping and/or peak-filling. Here we develop the ncGTW method to first detect misaligned features and then to realigned them. One unique advantage of ncGTW is to incorporate the structural information into multiple alignment, specifically concerning neighboring and distant samples with different RT drift patterns or degrees. To the best of our knowledge, most existing alignment methods have overlooked structural information related experiment design and batch duration. Moreover, the novel design of a reference-free multiple alignment strategy and utility of individualized warping functions across m/z bins all contributed to the expected superior performance of ncGTW.

Specifically designed to address the problem of misalignments complementary to existing alignment software tools, our proposed ncGTW method focuses on correcting only those misaligned features. Toward this objective with high efficiency, the ncGTW package includes a unique functional module that specifically aims to detect misaligned features. The outperformance of ncGTW is particularly attractive when processing large-scale datasets consisting of hundreds or thousands of samples, because the RT drifts between distant samples would be significant and warping functions over different m/z bins would be different (**Fig. 1**), due to long period of data acquisition time. Expectedly, explicit incorporation of RT structural information by ncGTW method helps achieving accurate realignments on the misaligned features. While we have only demonstrated ncGTW as a plug-in package to XCMS, in fact, two major functions of ncGTW tool can serve as a plug-in jointly or independently to other alignment tools as well, thus have broad applicability, even not limited to LC-MS datatype.

Regarding peak distortion problem associated with ncGTW, there are at least two potentially effective solutions. First, the peak information readily provided by XCMS can be utilized by ncGTW to correct peak distortion as we have discussed in section 2.4. Second, parameter settings in ncGTW can be adjusted or optimized to reducing the likelihood of peak distortion. Note that the current ncGTW tool package already includes a peak distortion correction module, and our experiments have also shown that interim peak-distortion correction can help optimize ncGTW parameter settings that will in turn reduce the likelihood of peak distortion.

In our misalignment detection step in section 3.4, we have observed that the false positive rate in the Rotterdam dataset is much higher than theoretical threshold of 0.05. By a closer look at the peak detection results, we found that many peaks were actually missed by XCMS, mainly due to significant yet irregular signal intensity fluctuating over the course of data acquisition as shown in **Fig. S11** with all samples. Considering such relatively higher false positive rate does not create significant computational burden on ncGTW yet may be uncontrollable, we have opted to first ‘accept’ these false positives and then screen them out at later stage.

## ACKNOWLEDGMENTS

This work was funded in part by the National Institutes of Health under Grants HL111362-05A1 and HL133932.

## AUTHOR CONTRIBUTIONS

C.T.W. developed ncGTW framework and wrote manuscript; D.M.H. designed the project with datasets; C.T.W. and Y.W. implemented ncGTW and GTW algorithms and performed real data analysis; Y.W. and G.Y. revised methodology and edited the manuscript; T. E. and I.K. provided expertise support.

## REFERENCES

Arnold, B.C., Balakrishnan, N. and Nagaraja, H.N. A first course in order statistics. Siam; 1992.

Bild, D.E., et al. Multi-Ethnic Study of Atherosclerosis: objectives and design. Am J Epidemiol 2002;156(9):871–881.

Christin, C., et al. Optimized time alignment algorithm for LC− MS data: correlation optimized warping using component detection algorithm-selected mass chromatograms. Analytical chemistry 2008;80(18):7012–7021.

Goldberg, A.V., et al. Maximum flows by incremental breadth-first search. In, European Symposium on Algorithms. Springer; 2011. p. 457–468.

Hofman, A., et al. The Rotterdam Study: 2014 objectives and design update. Eur J Epidemiol 2013;28(11):889–926.

Jiang, W., et al. Comparisons of five algorithms for chromatogram alignment. Chromatographia 2013;76(17-18):1067–1078.

Katajamaa, M. and Oresic, M. Processing methods for differential analysis of LC/MS profile data. BMC Bioinformatics 2005;6:179.

Lewis, M.R., et al. Development and Application of Ultra-Performance Liquid Chromatography-TOF MS for Precision Large Scale Urinary Metabolic Phenotyping. Anal Chem 2016;88(18):9004–9013.

Listgarten, J., et al. Multiple alignment of continuous time series. In, Advances in neural information processing systems. 2005. p. 817–824.

Lu, W., Bennett, B.D. and Rabinowitz, J.D. Analytical strategies for LC-MS-based targeted metabolomics. J Chromatogr B Analyt Technol Biomed Life Sci 2008;871(2):236–242.

Lu, X., et al. LC-MS-based metabonomics analysis. J Chromatogr B Analyt Technol Biomed Life Sci 2008;866(1-2):64–76.

Matuszewski, B., Constanzer, M. and Chavez-Eng, C. Matrix effect in quantitative LC/MS/MS analyses of biological fluids: a method for determination of finasteride in human plasma at picogram per milliliter concentrations. Analytical Chemistry 1998;70(5):882–889.

Mueller, L.N., et al. SuperHirn - a novel tool for high resolution LC-MS-based peptide/protein profiling. Proteomics 2007;7(19):3470–3480.

Petitjean, F., Ketterlin, A. and Gançarski, P. A global averaging method for dynamic time warping, with applications to clustering. Pattern Recognition 2011;44(3):678–693.

Prakash, A., et al. Signal maps for mass spectrometry-based comparative proteomics. Mol Cell Proteomics 2006;5(3):423–432.

Prince, J.T. and Marcotte, E.M. Chromatographic alignment of ESI-LC-MS proteomics data sets by ordered bijective interpolated warping. Analytical Chemistry 2006;78(17):6140–6152.

Sakoe, H., et al. Dynamic programming algorithm optimization for spoken word recognition. Readings in speech recognition 1990;159:224.

Smith, C.A., et al. XCMS: processing mass spectrometry data for metabolite profiling using nonlinear peak alignment, matching, and identification. Analytical chemistry 2006;78(3):779–787.

Smith, R., Ventura, D. and Prince, J.T. LC-MS alignment in theory and practice: a comprehensive algorithmic review. Brief Bioinform 2015;16(1):104–117.

Theodoridis, G., Gika, H.G. and Wilson, I.D. LC-MS-based methodology for global metabolite profiling in metabonomics/metabolomics. TrAC Trends in Analytical Chemistry 2008;27(3):251–260.

Tsugawa, H., et al. MS-DIAL: data-independent MS/MS deconvolution for comprehensive metabolome analysis. Nat Methods 2015;12(6):523–526.

Vinaixa, M., et al. A Guideline to Univariate Statistical Analysis for LC/MS-Based Untargeted Metabolomics-Derived Data. Metabolites 2012;2(4):775–795.

Wang, Y., et al. Graphical time warping for joint alignment of multiple curves. In, Advances in Neural Information Processing Systems. 2016. p. 3648–3656.

Zhang, X., et al. Data pre-processing in liquid chromatography-mass spectrometry-based proteomics. Bioinformatics 2005;21(21):4054–4059.

