## Supplementary Information for "Alignment of LC-MS Profiles by Neighbor-wise Compound-specific Graphical Time Warping with Misalignment Detection"

### Detection of Misaligned Features

We define a test statistic and derive the $p$-value for each initially-aligned feature from XCMS. There are total $N$ samples, and for a feature detected in $n$ samples, we denote the indices (run-order) of these samples as a set $\{l_1, l_2, \ldots, l_n\}$, where $1 \le l_i \le N$ and $l_1 < l_2 < \cdots < l_n$. For this feature, we define the test statistic

$$t = \max_{1 \le i, j \le n} \left| l_i - l_j \right|, \qquad (S1)$$

which is the same as the *range* of the $n$ indices. It is clear that $n - 1 \le t \le N - 1$. Under the null hypothesis, these $n$ samples are randomly drawn from the $N$ samples. Therefore, each index in this feature can be viewed as a random variable and we assume it follows a discrete uniform distribution. Our goal is to find the distribution of the random variable which is obtained by subtracting the smallest of these $n$ random variable from the largest one. To achieve this, we sort these $n$ random variables from small to large. The distribution of each *sorted* random variable is no longer uniform and can be calculated using order statistics (Arnold, et al., 1992). For $l_i$, the $i$th smallest index in the feature, the probability mass function (pmf) is

$$f_{l_i}(k) = \frac{\binom{k-1}{i-1} \binom{N-k}{n-i}}{\binom{N}{n}}, i \le k \le N - n + i, \qquad (S2)$$

and the joint pmf of $l_i$ and $l_j$ is given by

$$f_{l_i, l_j}(k, l) = \frac{\binom{k-1}{i-1} \binom{l-k-1}{j-i-1} \binom{N-l}{n-j}}{\binom{N}{n}}, i \le k < l \le N - n + j, l - k \ge j - i. \qquad (S3)$$

To obtain the null distribution of the test statistics, we let $i = 1, j = n, l = k + t$. Thus, equation (S3) becomes

$$f_{l_1, l_n}(k, k+t) = \frac{\binom{t-1}{n-2}}{\binom{N}{n}}, 1 \le k < k + t \le N, t \ge n - 1. \quad (S4)$$

Therefore, the pmf of $t$ is

$$f_T(t) = \sum_{k=1}^{N-t} f_{l_1, l_n}(k, k+t) = (N-t)\frac{\binom{t-1}{n-2}}{\binom{N}{n}}, n - 1 \le t \le N - 1. \quad (S5)$$

For an observed feature, we can calculate its *p*-value:

$$p\text{-}value = \Pr\{t \le t_{obs}\} = \sum_{t=n-1}^{t_{obs}} (N-t)\frac{\binom{t-1}{n-2}}{\binom{N}{n}}, \quad (S6)$$

where $t_{obs}$ is the test statistic calculated in that feature (Connor, 1969). The summation is over all $t$s that are smaller than $t_{obs}$, which reflects the assumption that the more concentrated the indices are, the more likely misalignment exists. The smaller the p-value, the more unlikely the observed feature follows the null distribution, and the more likely misalignment occurs.

### Detailed on the alignment module of ncGTW

ncGTW contains a multiple alignment module that is designed to flexibly model and incorporate any structural information among samples, as long as their relationship can be represented as a weighted graph. This module aims to find a set of warping functions, through which each sample can be aligned to a reference. The main challenge in multiple alignment is the lack of a priori reference. Indeed, various strategies have been proposed to select a reference sample or estimate a reference using all samples. However, when the samples have complex patterns such as missing signals at some time points, or contain significant noise, no single sample merits a good reference while the estimation of an ensemble reference requires a set of pre-aligned samples.

Without picking a certain reference, ncGTW extracts the needed information from all possible sample pairs. To deal with missing signals and noise, ncGTW borrows the idea from graphical time warping (GTW) (Wang, et al., 2016) to incorporate the structure information in the dataset, so that the estimation of the pairwise warping functions becomes more accurate. The flowchart of ncGTW is shown in **Fig. 4** in the main article. We first find all pairwise warping functions with the structural prior knowledge as constraints. Then we set all pairwise warping functions as constraints to estimate the final warping functions (the warping functions for samples aligning to a reference) of all samples. These two subproblems are formulated and solved with the framework of network flow algorithms that can produce a global and efficient solution.

#### Problem modeling and methods

Given $N$ samples (curves) $\{x_1, \ldots, x_N\}$, multiple alignment problem aims to find a set of warping functions $\{\Phi_{i,c}\}, i \in \{1 \ldots N\}$, through which each sample can be aligned to a reference $x_c$. For the sake of clarity, all samples have the same number of points $P$. The subscript "$i, c$" means this function maps a set of points $\{x_{ip}\}, p \in \{1 \dots P\}$ in curve $x_i$ to a corresponding set of points in curve $x_c$. That is, for any point $x_{ip}$ in $x_i$, $\Phi_{i,c}$ can always map $x_{ip}$ to at least one point in $x_c$, where $p$ is from 1 to $P$.

**Definition 1 – valid warping function**

A valid warping function for the pair of curves $(x_i, x_j)$ is a set of integer pairs $\Phi_{i,j} = \{(p, q)\}$, such that the following conditions are satisfied: (a) boundary conditions: $(1,1) \in \Phi_{i,j}$ and $(P, P) \in \Phi_{i,j}$; (b) continuity and monotonicity conditions: if $(p, q) \in \Phi_{i,j}$, then $(p-1, q) \in \Phi_{i,j}$ or $(p, q-1) \in \Phi_{i,j}$ or $(p-1, q-1) \in \Phi_{i,j}$. An example is shown in **Fig. S1a**.

**Definition 2 – inverse of a valid warping function**

Given a valid warping function $\Phi_{i,j} = \{(p, q)\}$, the inverse of $\Phi_{i,j}$ is ${\Phi_{i,j}}^{-1} = \{(q, p)\} = \Phi_{j,i}$

**Definition 3 – alignment cost**

For any given valid warping function $\Phi_{i,j}$ and its corresponding pair of curves $(x_i, x_j)$, the associated alignment cost is defined as follows:

$$\mathrm{cost}(\Phi_{i,j}) = \sum_{(p,q)\in\Phi_{i,j}} g(\mathrm{x}_{ip} - \mathrm{x}_{\mathrm{j}q}), \qquad (S7)$$

where $g(\mathrm{x}_{ip} - \mathrm{x}_{\mathrm{j}q})$ is any nonnegative function which compute the distance between $\mathrm{x}_{ip}$ and $\mathrm{x}_{\mathrm{j}q}$.

#### Stage 1: jointly aligning all pairs with the structural prior incorporated

To estimate $\{\Phi_{i,j}\}$ jointly, ncGTW considers all possible sample pairs. For each pair, one sample is set as the reference. In other words, in the first stage, ncGTW tries to align each sample to the other samples. For $N$ samples, the number of alignment pairs is $N(N-1)$. However, $\Phi_{i,j}$ is the inverse of $\Phi_{j,i}$, so only $N(N-1)/2$ pairs need to be considered in the real implement. That is, only $N(N-1)/2$ pairwise warping functions are needed.

In order to incorporate the structure information, as GTW, we need to convert the given structure information in the dataset into the warping function neighborhood information. Suppose $(x_i, x_j)$ are neighbors by structure information, we consider the pair of warping functions, $(\Phi_{i,k}, \Phi_{j,k})$, as well as $(\Phi_{k,i}, \Phi_{k,j})$, as neighbors respectively for all $k$. Also, if $(x_i, x_j)$ are neighbors and $(x_k, x_l)$ are also neighbors respectively, we can consider the pair of $\Phi_{i,k}$ and $\Phi_{j,l}$ are neighbors. **Fig. S1c** gives an example of the warping function neighborhood information of five samples. The run orders of the five samples are exactly 1, 2, 3, 4, and 5. Thus, these five samples are expected to have a pattern of continuous changing, so $(x_1, x_2)$, $(x_2, x_3)$, $(x_3, x_4)$, and $(x_4, x_5)$ are considered as neighbors respectively. More examples about the structures are shown in the experiments section.

**Definition 4 – neighboring warping functions**

Suppose the neighboring structure for a set of $M$ valid warping functions is given by the graph $G_{struct} = \{V_s, E_s\}$, where $V_s$ is the set of nodes, with each node corresponding to a warping function, and $E_s$ is the set of undirected edges between nodes. If $v_{ij}, v_{kl} \in V_s$ and $(v_{ij}, v_{kl}) \in E_s$, we call $\Phi_{i,j}$ and $\Phi_{k,l}$ neighbors, denoted by $((i,j),(k,l)) \in Neib$.

As an improvement of GTW, ncGTW also adapts the idea of dynamic time warping (DTW). DTW aligns two curves $x_i$ and $x_j$ by finding a warping function $\Phi_{i,j}$ that minimize the alignment cost (S7). The correspondence represented by $\Phi_{i,j}$ can be visualized as a path in a DTW grid, from bottom left to top right, and the weight of each edge is decided by the distance function $g(\cdot)$ with all possible point pairs $(x_{ip}, x_{jq})$, where $x_{ip} \in x_i$ and $x_{jq} \in x_j$. DTW estimates $\Phi$ in the

DTW grid using dynamic programming and various additional constraints can be employed, such as the direction of the path.

**Definition 5 – DTW grid for a single pair of curves**

For each pair of curves, consistent with the cost function (1), there is an induced directed planar graph (Korte, et al., 2012), $G_{ij} \coloneqq \{V_{ij}, E_{ij}\}, 1 \leq i < j \leq N$, where $V_{ij}$ and $E_{ij}$ are the nodes and directed edges respectively. Each point pair $\left(x_{ip}, x_{jq}\right)$ is corresponding to a node $V_{ij,pq} \in V_{ij}$, where $1 \leq p, q \leq P$. The weight of $\left(V_{ij,p_1q_1}, V_{ij,p_2q_2}\right) \in E_{ij}$ is the distance between the two points $(x_{ip_1}, x_{jq_1})$, measured by $g\left(x_{ip_1} - x_{jq_1}\right)$. An example is shown in **Fig. S1a**. Any directed path from the bottom-left corner to the upper-right corner is corresponding to a valid warping function $\Phi_{i,j}$.

Once the structure for warping functions is obtained, the joint alignment of all pairs of samples can be readily solved by the recently developed model – Graphical Time Warping (GTW). When we jointly align multiple pairs of curves with structure information, our goal is to minimize both the overall alignment cost and the distance between neighboring warping functions.

**Definition 6 – distance between two valid warping functions**

For any two given valid warping functions $\Phi_{i,j}$ and $\Phi_{k,l}$, the distance between them is defined as follows:

$$\operatorname{dist}\left(\Phi_{i,j}, \Phi_{k,l}\right) = \frac{1}{2} \sum_{1 \leq n \leq P} \left| \max_{(p_i,n) \in \Phi_{i,j}} p_i - \max_{(p_k,n) \in \Phi_{k,l}} p_k \right| + \left| \min_{(p_i,n) \in \Phi_{i,j}} p_i - \min_{(p_k,n) \in \Phi_{k,l}} p_k \right|, \qquad (S8)$$

which is equivalent to the area of the region bounded by the two corresponding paths in a DTW grid as shown in **Fig. S1a**.

Mathematically, to balance the alignment cost (S7) and the distance between warping functions (S8), we want to solve the following GTW problem:

$$\min_{\Phi} f\,(\Phi) = \min_{\Phi=\{\Phi_{i,j}|1\le i<j\le N\}} \sum_{1\le i<j\le N} \mathrm{cost}(\Phi_{i,j}) + \kappa_1 \sum_{((\mathrm{i,j}),(\mathrm{k,l}))\in \mathrm{Neib}} \mathrm{dist}(\Phi_{i,j}, \Phi_{k,l}), \qquad (S9)$$

where $\kappa_1$ is the parameter which balance the two terms. The first term is the overall alignment cost of the warping functions. The second term could be considered as the sum of the "dissimilarity" of the neighboring warping functions. Again, the neighboring warping functions should be similar. In the other words, their distance should be small.

There are three major steps in to solve the GTW problem. Firstly, GTW transforms the DTW grid for each pair of curves to an equivalent minimum cut problem as a DTW graph. Supposing there are *M* pairs of curves, where each pair contains two curves to be aligned with each other, then we will have *M* DTW grids, of which there are also *M* DTW graphs. Secondly, GTW adds in extra edges between DTW graphs, if two pairs of curves are considered as neighbors. Note that if edges are to be added between two DTW graphs, all the corresponding vertices in the two graphs need to be connected. Thus, the *M* separate DTW graphs become an extended graph (GTW graph). Thirdly, GTW proved that the joint alignment of multiple pairs with smoothness constraints imposed could be formulated as a minimum cut problem in the GTW graph. Hence, efficient network flow algorithms can be used to find a global solution.

**Definition 7 – DTW graph**

Define $G'_{ij} \coloneqq \{V'_{ij}, E'_{ij}\}$ as the DTW graph of the DTW grid $G_{ij}$, where nodes $V'_{ij}$ are all faces of $G_{ij}$. That is, for each $V_{ij,pq} \in V_{ij}$, where $2 \le p \le P-1$ and $1 \le q \le P-1$, there are two corresponding nodes $V'_{ij,pq+}$ and $V'_{ij,pq-}$; for each $V_{ij,pq}$ where $p = 1$ and $1 \le q \le P-1$, there is one corresponding nodes $V'_{ij,pq+}$; for each $V_{ij,pq}$ where $p = P$ and $1 \leq q \leq P-1$, there is one corresponding nodes $V'_{ij,pq-}$. For each $e \in E_{ij}$, we have a new edge $e' \in E'_{ij}$ connecting the faces from the right side of $e$ to the left side. This edge is directed (with positive direction by convention). The edge weights are the same as for the primal graph $G_{ij}$. An example is shown in **Fig. S1b**.

**Definition 8 – GTW graph**

The GTW graph $G_{gtw} \coloneqq \{V_{gtw}, E_{gtw}\}$ is defined as the *integrated graph* of all DTW graphs $\{G'_{ij} | 1 \leq i < j \leq N\}$ with the integration guided by the neighborhood of warping functions, such that $V_{gtw} = \{V'_{ij} | 1 \leq i < j \leq N\}$ and

$$E_{gtw} = E'_{ij} \big| 1 \leq i < j \leq N \cup \{(V'_{ij,pq+}, V'_{kl,pq+}), (V'_{ij,pq-}, V'_{kl,pq-})\} \big| ((\mathrm{i},\mathrm{j}), (\mathrm{k},\mathrm{l})) \in Neib.$$

All newly introduced edges $(V'_{ij,pq+}, V'_{kl,pq+})$ and $(V'_{ij,pq-}, V'_{kl,pq-})$ are bi-directional with capacity $\lambda_1$ as shown in **Fig. S1d**. An example of a GTW graph with two pairs of curves is shown in **Fig. S1e**.

**Definition 9** – **Labeling of the graph**

$L$ is a *labeling* of graph $G$ if it assigns each node in $G$ a binary label. $L$ can induce a cut set $C = \{(s,t) | L(s) \neq L(t), (s,t) \in E_G\}$ .The corresponding cut (or flow) is $cut(L) = cut(C) = \sum_{(s,t) \in C} weight(s,t)$, where $weight(s,t)$ is the weight on the edge between nodes $s$ and $t$.

Based on its construction, a labeling $L$ for the graph $G_{gtw}$ can be written as $L = \{L_{ij} | 1 \leq i < j \leq N\}$, where $L_{ij}$ is a labeling for the DTW graph $G'_{ij}$. Thus, we can express the minimum cut problem for the graph $G_{gtw}$ as:

$$\min_{L} g(L) = \min_{L:=\{L_{ij}|1\le i<j\le N\}} \sum_{1\le i<j\le N} cut(L_{ij}) + \lambda_1 \sum_{((i,j),(k,l))\in Neib} cut(L_{ij}, L_{kl}), \qquad (S10)$$

where $cut(L_{ij})$ is the cut of all edges for $G'_{ij}$ and $cut(L_{ij}, L_{kl})$ is the number of the cut edges between two neighboring dual graphs $G'_{ij}$ and $G'_{kl}$.

As proved in (Wang, et al., 2016), the GTW problem as stated in (S9) is equivalent to the minimum cut problem on the GTW graph $G_{gtw}$ if we set $\lambda_1 = 2\kappa_1$. Once we apply any existing network flow algorithm to solve this minimum cut problem, the GTW problem would be solved, and all pairwise warping functions are available.

#### Stage 2: Finding multiple alignment based on the constraint of pairwise alignments

In the stage 2 of ncGTW, the result from the stage 1, $\{\Phi_{i,j}\}$, is used to estimate $\{\Phi_{i,c}\}$, which are the final goal of multiple alignment problem. Here, $N$ warping functions need to be estimated. On the contrary to the stage 1, where neighboring information as constraints, in stage 2 the warping function $\{\Phi_{i,j}\}$ are set as constraints to estimate $\{\Phi_{i,c}\}$. For example, according to the warping function $\Phi_{i,j}$, point $x_{ip}$ in $x_i$ should be aligned to point $x_{jq}$ in $x_j$. Then, we expect that $x_{ip}$ and $x_{jq}$ should be aligned to the same position on the reference $x_c$. That is, we want the warping of $x_{ip}$ and $x_{jq}$ on the reference is *consistent*.

**Definition 10 – inconsistency between two sample points**

For any given integer pair $(p_i, p_j) \in \Phi_{i,j}$, where $p_i$ is the point index of $x_i$ and $p_j$ is the point index of $x_j$. The inconsistency between $x_{ip_i}$ and $x_{jp_j}$ is defined as follows:

$$\text{incons}\left(x_{ip_i}, x_{jp_j}; \Phi_{i,j}\right) = \left|\max_{(p_i,q_i)\in\Phi_{i,c}} q_i - \max_{(p_j,q_j)\in\Phi_{j,c}} q_j\right| + \left|\min_{(p_i,q_i)\in\Phi_{i,c}} q_i - \min_{(p_j,q_j)\in\Phi_{j,c}} q_j\right|,$$

which could be considered as the distance between two continuous integer sets, as shown in **Fig. S2a**. The inconsistency quantifies how much the alignment deviates from our expectation. If the inconsistency is zero, we call these two points are consistent.

**Definition 11 – inconsistency between two warping functions**

For any given valid warping function $\Phi_{i,j}$, the inconsistency between two warping functions $\Phi_{i,c}$ and $\Phi_{j,c}$ is defined as follows:

$$\mathrm{incons}\left(\Phi_{i,c}, \Phi_{j,c}; \Phi_{i,j}\right) = \sum_{(p_i,p_j)\in\Phi_{i,j}} \left| \max_{(p_i,q_i)\in\Phi_{i,c}} q_i - \max_{(p_j,q_j)\in\Phi_{j,c}} q_j \right| + \left| \min_{(p_i,q_i)\in\Phi_{i,c}} q_i - \min_{(p_j,q_j)\in\Phi_{j,c}} q_j \right|, (S11)$$

which is equivalent to the sum of the inconsistency between two sample points $x_{ip_i}$ and $x_{jp_j}$, where $(p_i, p_j) \in \Phi_{i,j}$. Likewise, if the inconsistency between these two warping functions is zero, then the two warping functions are consistent.

Intuitively speaking, stage 2 tries to identify the final warping functions from all pairwise ones, with the constraint that the sum of inconsistency between all warping function pairs as low as possible. However, due to the noise and other factors, in the real data, there are always some contradictions among the pairwise warping functions. For example, from $\Phi_{i,j}$, we know that $x_{ip}$ and $x_{jq}$ are aligned together, and from $\Phi_{j,k}$, we know that $x_{jq}$ and $x_{kr}$ are aligned together, but from $\Phi_{k,i}$, we know that $x_{kr}$ and $x_{io}$ not $x_{ip}$ are aligned together. This kind of contradictions will make the alignment tend to be the trivial "all to one" mapping, if we want to minimize the inconsistency. To solve this problem, here we introduce another constraint that makes the warping functions tend to choose "one to one" mapping, to avoid the trivial mapping. In the real implement, we redesign the weight of the edges in the DTW graphs. Thus, the exact values of neither $x_i$ nor

$x_c$ do no matter in this stage. In the other words, the weights of the edges on the DTW graph of $x_i$ and $x_c$ is not based on the points of $x_i$ and $x_c$. This property is the reason why our approach does not rely on the selection or estimate of the reference curve. Therefore, $x_c$ is called a "virtual reference".

**Definition 12 – nondiagonality of a warping function**

For any given valid warping function $\Phi_{i,c}$, the associated nondiagonality is defined as follows:

$$\text{nondiag}\left(\Phi_{i,c}\right) = \sum_{(p,q)\in\Phi_{i,c}} \mathbb{1}\left((p-1,q) \in \Phi_{i,c} \vee (p,q-1) \in \Phi_{i,c}\right), \qquad (S12)$$

where $\mathbb{1}(\cdot)$ is the indicator function. This definition is as same as the total number of vertical and horizontal edges in the corresponding path. The smallest value of nondiagonality is zero (one-to-one mapping, the diagonal line). An example of nondiagonality is shown in **Fig. S2a**.

In order to obtain the consensus final alignment from pairwise warping functions, we design ncGTW problem which tries to balance the nondiagonality (S12) and inconsistency (S11) among final warping functions:

$$\min_{\Phi} f\,(\Phi) = \min_{\Phi=\{\Phi_{i,c}|1\le i\le N\}} \sum_{1\le i\le N} \text{nondiag}\left(\Phi_{i,c}\right) + \kappa_2 \sum_{1\le i<j\le N} \text{incons}\left(\Phi_{i,c}, \Phi_{j,c}; \Phi_{i,j}\right), \qquad (S13)$$

where the first term relates to the warping path for each final warping function, and the second term is the constraints between each warping function pair. The relation between these two terms is similar to the two terms in the GTW problem (S9). Thus, as same as GTW, an ncGTW problem could also be transformed to an ncGTW graph, and solved by any network flow algorithm.

**Definition 13 – ncGTW graph**

The ncGTW graph $G_{ncgtw} \coloneqq \{V_{ncgtw}, E_{ncgtw}\}$ is defined as the *integrated graph* of all \DTW graphs $\{G'_{ic} | 1 \le i \le N\}$ with the integration guided by the pairwise warping functions, such that $V_{ncgtw} = \{V'_{ic} | 1 \le i \le N\}$ and $E_{ncgtw} = \{E'_{ic} | 1 \le i \le N \cup \left(V'_{ic,p_iq+}, V'_{jc,p_jq+}\right) | (p_i, p_j) \in \Phi_{i,j} \cup \left(V'_{ic,p_iq-}, V'_{jc,p_jq-}\right) | (p_i, p_j) \in \Phi_{i,j}\}$. That is, all newly introduced edges are guided by $\Phi_{i,j}$ as shown in **Fig. S2b**. Also, all these new edges are bi-directional with capacity $\lambda_2$. For the edges in $V'_{ic}$, the capacity of edges corresponding to vertical or horizontal path is one, and zero for edges corresponding to diagonal path, so that the nondiagonality of the warping function $\Phi_{i,c}$ is equivalent to the cost of the warping path on the corresponding DTW grid $G_{ic}$.

Like GTW problem, the ncGTW problem as stated in equation (S13) is equivalent to the minimum cut problem on the ncGTW graph $G_{ncgtw}$ if we set $\lambda_2 = 2\kappa_2$. Moreover, a labeling $L$ for the graph $G_{ncgtw}$ can be written as $L = \{L_{ic} | 1 \le i \le N\}$, where $L_{ic}$ is a labeling for the DTW graph $G'_{ic}$. Therefore, we can express the minimum cut problem for the graph $G_{ncgtw}$ as:

$$\min_L g(L) = \min_{L \coloneqq \{L_{ic} | 1 \le i \le N\}} \sum_{1 \le i \le N} cut(L_{ic}) + \lambda_2 \sum_{1 \le i < j \le N} cut\left(L_{ic}, L_{jc}\right), \qquad (S14)$$

where $cut(L_{ic})$ is the cut of all edges for $G'_{ic}$ and $cut\left(L_{ic}, L_{jc}\right)$ is the number of the cut edges between two neighboring dual graphs $G'_{ic}$ and $G'_{jc}$.

Again, after applying any existing network flow algorithm, the ncGTW problem would be solved, and all final warping functions $\{\Phi_{i,c}\}$ are available.

#### Local or global optimum

In the stage 1 of ncGTW, we claimed that we could obtain the global optimum of a GTW problem. That is, we can solve the maximum flow problem from the GTW graph with the global optimum.

One should notice that this global optimum is not the global optimum of the multiple alignment problem. It is possible that the global optimum of GTW is just the local optimum of multiple alignment. Though ncGTW could not guarantee to find the global optimum of the multiple alignment problem as predicted by the fact that multiple alignment problem is an NP-hard problem, ncGTW still has a superior advantage. In the different fields, the ways of evaluation of the result of multiple alignment are very different. However, ncGTW can adjust the weights of the additional edges according to the evaluation method. Thus, with the structure information, ncGTW can approach to the global optimum of the multiple alignment better than other methods, with great flexibility to various evaluation criteria.

### Experiments

We evaluate the performance of ncGTW on both simulation and real datasets. Any neighborhood structure among samples can be incorporated into ncGTW as long as the structure can be represented as a graph. Although weights on edges can also be naturally integrated into ncGTW, prior knowledge on weights is very application-dependent and thus we assume the neighborhood structure has no weight. In this set of experiments, we test three different structures: line, block, and uniform (non-informative). Three peer methods, DBA, CPM, and GTW were selected for comparison due to their representativeness. DBA iteratively computes barycenter and aligns all samples to the barycenter (Petitjean, et al., 2011). CPM is a classical probability model method and uses the hidden Markov model to learn the prototype function (Listgarten, et al., 2005). For GTW, since it needs a reference and GTW methodology does not provide a way to select a reference, to apply GTW to the multiple alignment problem we use all samples as reference one by one and take the average of all scores. For a visual demonstration, we choose the most informative one. In the following experiments, one can see that a bad reference may ruin the alignment of GTW. Also, other flaws of directly applying GTW on multiple alignment are demonstrated.

Both classical measurements and visual assessment were used to evaluate performance. We adopt two quantitative criteria that are frequently used in the literature. They are mean correlation coefficient (MCC) and simplicity (SP) (Jiang, et al., 2013). After alignment, the correlation among samples is expected to increase. Thus, the mean of the correlation of all sample pairs can be considered as a quantification of the alignment quality. The definition of simplicity here is the sum of the fourth power of all singular values, where the sum of all singular values is normalized to be one. The singular values are computed based on the data matrix, where each row represents a sample. The idea is that if it is a good alignment, the first singular value should dominate others. Hence, larger simplicity means better alignment. A note is that the largest possible value for simplicity is one.

For the simulation data, since we have the ground truth, we test each peak separately so that we can know the alignment quality of each peak. For the real data, the numerical evaluations for each peak are not applicable since we do not have the ground truth.

#### Case study on line structure

When samples change gradually according to a variable such as time, we call it to have a line structure, which can be converted to triangles (**Fig. S3b**) as an input structure between pairwise alignments. To evaluate the performance of different methods, we first designed a simulation dataset containing five samples. The first sample is shown in **Fig. S4a** and all five samples are plotted in **Fig. S4b**. Note that all the samples contain three peaks, except the fourth sample (purple dash line, the third peak of which is missing. This phenomenon of missing peaks occurs frequently in real applications.

**Fig. S4c-f** shows the alignment result of DBA, CPM, GTW, and ncGTW, and **Table S1** shows the evaluation scores of each peak. As a ground truth, the two peaks pointed by black arrows belong to the first and second group, respectively. The MCC and SP of all peaks of all methods are improved, compared with the original curve. However, DBA wrongly aligned the peaks in sample 4, which leads to bad scores of MCC and SP. CPM aligned all the peaks correctly, so it got not bad scores for all three peak groups in three evaluation. In **Fig. S4e**, the reference of GTW is the fourth sample. Since the fourth sample lacks the third peak, we can see from the figure that the third peaks of all samples are not aligned well. This is an example of that the reference may have a huge effect on alignment.  Moreover, the variance of the fourth sample is the largest one. In general, when needing a reference, many existing methods posit that the sample with the largest variance should be selected, so here we also show that reference selecting is a hard problem. Even with averaging, the scores of peak 3 of GTW are still much worse than other methods. Our newly proposed method ncGTW produced accurate alignment as evidenced by both visual assessment and quantitative scores. One may notice that the MCC of peak 3 are all smaller than 0.6, which is due to the missing peak in the fourth sample.

#### Case study on block structure

Block structure means there are several blocks formed by samples in the dataset. Within each block, the shifts are small. Between blocks, the shift is larger. Suppose we have ten samples in a dataset, and every five samples form a block. **Fig. S5** shows how to connect these pairs. We can separate these pairs into three types. The first type is the alignment within the first block. The second type is between the two blocks. The third type is within the second block. Only the same type of neighbors will be connected. To test this structure, we used a ten-sample dataset, and every five samples form a block. **Fig. S6a** shows the first sample, which contains three peaks. **Fig. S6b** shows the eighth sample, of which the third peak is missing. **Fig. S6c-d** show the two blocks in the dataset. **Fig. S6e** shows all the samples.

**Fig. S6f- i** show the results of DBA, CPM, GTW, and ncGTW, and **Table S2** shows the peak scores for each method. DBA wrongly aligned the two peaks in the eighth sample, due to the same reason as in the previous section. CPM has the worst performance since CPM separated all the peaks into four groups, not three. The reason may be that in **Fig. S6e**, it somehow shows four peak groups, and in each group, there are at least five peaks. Thus, CPM considered there are four peak groups, not three. The reference of GTW is the tenth sample, which contains three peaks. With a good reference, in **Fig. S6h**, GTW have similar performance as good as ncGTW. However, since there is a missing peak in the eighth sample, after averaging, the MCC of peak 3 is a little bit lower than ncGTW. Again, the MCC of peak 3 is not close to one for all methods, and this is also due to the missing peak in the eighth sample.

#### Case study on non-informative structure

Sometimes we may know nothing about the structure of the dataset. In this section, we demonstrate the experiments on the dataset without any structure information. For simulation, here we consider a ten-sample dataset. Without structure information, in the first stage of ncGTW, we add edges between the pairs that have the same aligning sample or reference sample. For example, for $G'_{1,2}$, we will connect it to $G'_{1,i}$, where $i$ is from 3 to 10. Likewise, for $G'_{2,1}$, we will connect it to $G'_{i,1}$, where $i$ is from 3 to 10. In this 10 simulated samples dataset, there are two samples without peak 1, another two without peak 2, and still another one without peak 3. Thus, there are only four samples with all three peaks. **Fig. S7a** shows a sample without peak 2. **Fig. S7b** shows all ten samples. **Fig. S7c-f** show the results of DBA, CPM, GTW, and ncGTW, and **Table S3** shows the peak scores for each method. DBA again wrongly aligned the sample shown in **Fig. S7a** (pointed by arrows). CPM aligned all the peaks well but with some small drifts in some samples and distortions, so the scores are not as good as ncGTW, but still better then DBA. With the sample shown in **Fig. S7a** as reference, GTW again misaligned the peak 2 group as shown in **Fig. S7e**. Since there are six samples with a peak missing, the average scores of GTW are all relatively low. Even without any structure information, ncGTW have the best performance.

#### Case study on real LC-MS dataset

In this section, we conducted tests on a real liquid chromatography-mass spectrometry (LC-MS) experiments. First, ten samples were selected and ordered by the time as they were assayed. Assuming the properties of the equipment were gradually changing, we impose a line structure among these samples. **Fig. S8a** shows one sample from the ten samples. Clearly, there are nine peaks in the sample. **Fig. S8b** shows all ten samples together. Note that the intensity of the corresponding peaks between samples are very different, and some peaks are missing.

**Fig. S8c-f** show the results of DBA, CPM, GTW, and ncGTW. DBA wrongly aligned the peak pointed by the arrow to the third group of peaks (should be aligned to the second group). The reason is at some steps, DTW aligned a peak to a peak with similar intensity, but these two peaks are not in the same group. DTW based methods have the tendency to align the peaks with similar intensities together. CPM also aligned the same peak wrongly to the third group. Similar to DBA, the intensity of the peak is so strong that CPM decided to align this peak to the group with a higher intensity. The reference sample of GTW is just the one shown in **Fig. S8a** and GTW aligned the arrow-pointed peak correctly, while the fourth and fifth peak groups were misaligned. Again, ncGTW aligned all the peaks well, so that the nine peak groups are clearly separated and none of them is clumped together.

Next, we chose 20 samples from two batches. Among the first ten samples, the retention time drift is small, also for the rest ten samples. However, between the first and the second blocks (batches), the drift is larger. Fig. **S9a-b** show an example of each block, and there are three peaks for each two sample. However, the third peak of most samples is missing. **Fig. S9c-d** show the first and second blocks in the dataset. There are three peak groups in both blocks. However, for the third peak group, only two samples in the first block and only three samples in the second block have the peak. To see other peaks clearer, in **Fig. S9e**, we show all the samples but change the scale of the y-axis.

**Fig. S9f-i** show the results of DBA, CPM, GTW, and ncGTW. DBA aligned more than ten peaks to the third peak group but should be five, with serious distortions. CPM also aligned wrongly the third peak group, since there are too many missing peaks in the third peak group. The reference of GTW is the first sample of the first block, as shown in **Fig. S9a**. Though there is no missing peak in the reference, GTW still tends to align the peaks with similar intensity together. Thus, the third peak group has more than five peaks, also with the distortion of peak shape. Only ncGTW aligned correctly the five peaks to the third peak group. This example demonstrated the significant improvement of ncGTW compared to GTW. We can see the advantage that ncGTW does not need a certain reference, and there is no distortion after alignment.

Next, to test ncGTW on data with no structure (or in the scenario that we are not sure about its structure), we use the same ten LC-MS samples in **Fig. S8b** but without the structural information. That is, we will add edges between the pairs with the same aligning sample or reference sample in the first stage of ncGTW.

**Fig. S8a-d** are as same as **Fig. S10a-d**, since DBA and CPM do not consider structural information already. As shown in **Fig. S10e-f**, unlike DBA and CPM, GTW and ncGTW still well aligned the pointed peak. However, without the structural information, GTW misaligned peak 3 and peak 4. Even without structural information, ncGTW accurately aligned all the peaks.

#### Discussion

Readers may be aware of the significant progress in aligning multiple DNA sequences. One may wonder why similar progress has not been seen in the general category of multiple alignment and why good ideas for DNA sequence alignment cannot be directly transferred to other applications.

In our view, this is not surprising because many effective approaches for DNA sequences explicitly or implicitly rely on the assumption of the existence of the evolution tree. Yet, the tree structure is very special and many applications cannot be described by a tree structure. Our ncGTW can be understood as an extension of tree structure to any graph structure. In addition, unlike methods designed for DNA sequences, our method models all samples simultaneously. Though we did not test our algorithm on DNA sequencing data, we expect to see favorable performance due to the integrative modeling nature.

Our approach is built on GTW, a recent extension of DTW to multiple pairs. GTW retains all desirable properties of DTW such as monotonicity of time shifts and polynomial efficient solution, and yet flexible models any graph-encoded structure among pairs. However, GTW cannot be directly applied to solve multiple alignment problem. GTW takes multiple pairs of samples as input and finds alignment for each pair with consistency between pairs considered. The problem considered in this paper takes multiple samples as input and aims to find consistent alignment among samples. Though we can manually specify one certain sample as a reference to construct pairs of samples for input of GTW, to our knowledge, there is no established method that can always help the user to identify which sample is the most suitable one as the reference. Moreover, it is likely that none of the samples contains enough information to serve as a good reference. That is, no matter we choose any sample as the reference, we can never obtain an accurate alignment. Thus, ncGTW is a significant improvement of GTW, since ncGTW can model all samples simultaneously and deal with multiple alignment problem without manually setting a reference.

The experiments clearly demonstrated the power of ncGTW. When structural information is available, it is anticipated to see increased accuracy of alignment due to the use of extra information. It is a little bit counterintuitive to observe that ncGTW still performed better when there was no informative structure. From hindsight, this is also expected because one can always borrow information from other samples if we use a Bayesian perspective and consider all samples forming a prior distribution. This phenomenon is also related to Stein’s paradox in the estimation theory (Efron and Morris, 1977).

### The Implementation of ncGTW on Large Dataset

For a large dataset, it is very time-consuming to simultaneously align all the samples for a profile-based alignment method. Take ncGTW for example, the node number of the graph in the first stage is propositional to the square of the sample number. When the sample number is huge, the graph becomes extremely big, and it may take hours or even days to solve the maximal flow no matter which algorithm is used. Thus, in the real implementation, we will split the whole dataset into several sub-datasets, and apply ncGTW to each small dataset. That is, in this way, the number of nodes in the graph in each small dataset will decrease significantly, comparing to the original graph. Also, since these small datasets are aligned independently, we can further reduce the computation time with parallel computing (depending on how much CPU cores you have in your computer). After the alignment of each small dataset is done, we build a “super-sample” for each small dataset and align these super-samples. With the warping functions within each small dataset and of the super-samples, we can obtain the final warping functions for each sample. We called this hierarchical alignment process as “two-layer ncGTW”. **Fig. S11** is the diagram of two-layer ncGTW. In the following sections, we will explain each layer in details.

#### The first layer of two-layer ncGTW

In the first layer, in the beginning, we need to decide how many sub-datasets we should split the original dataset into. For the computational efficiency, we recommend in each sub-dataset, there should be at least 10 samples. For example, if there are 500 samples in the original dataset, it is recommended to split the dataset into 10 sub-datasets (50 samples in each). Then, apply ncGTW on these sub-datasets independently. If the CPU cores the user have are more than 10, the computation time of this layer is equivalent to apply ncGTW on a 50-sample dataset once. After the alignment of each sub-dataset is done, we obtain the warping functions of all samples. With the warping functions, within each sub-dataset, the samples can be aligned to the same RT axis, and a "super-sample" will be generated by taking the average on the aligned samples. After the super sample of each sub-dataset is generated, we build a super-dataset with these super samples and send the super-dataset to the second layer with the warping functions of all samples.

##### The second layer of two-layer ncGTW

In the second layer, ncGTW is applied to the super-dataset to align the super-samples. Since the super-samples are similar to the common samples, we can directly apply ncGTW on them. Therefore, we can obtain a set of warping functions for super-samples. With the warping functions from the first layer (sample to super-sample) and the ones for super samples (super-sample to the final axis), all the samples can be aligned to the final axis, so that the alignment of all samples is done.

#### Supplementary Tables

| Peak1 | | |
|---|---|---|
| Scores / Methods | MCC | SP |
| Before alignment | 0.3071 | 0.4240 |
| DBA | 0.5683 | 0.5095 |
| CPM | 0.9697 | 0.9390 |
| GTW | 0.9988 | 0.9986 |
| ncGTW | 0.9999 | 0.9999 |
| **Peak2** | | |
| Scores / Methods | MCC | SP |
| Before alignment | 0.3072 | 0.4407 |
| DBA | 0.5671 | 0.5118 |
| CPM | 0.8943 | 0.8169 |
| GTW | 0.9999 | 0.9999 |
| ncGTW | 0.9999 | 0.9999 |
| **Peak3** | | |
| Scores / Methods | MCC | SP |
| Before alignment | 0.0848 | 0.3959 |
| DBA | 0.4912 | 0.9999 |
| CPM | 0.5042 | 0.9384 |
| GTW | 0.3986 | 0.8791 |
| ncGTW | 0.5099 | 0.9999 |

**Table S1.** Peak scores for four methods of line structure, where ‘Before alignment’ serves as the baseline, MCC and SP represent mean correlation coefficient and simplicity respectively, and the range of either score is between 0 and 1 with higher score indicating better performance.

| **Peak1** | | |
|---|---|---|
| Scores / Methods | MCC | SP |
| Before alignment | 0.1163 | 0.1937 |
| DBA | 0.7894 | 0.8326 |
| CPM | 0.3923 | 0.4603 |
| GTW | 0.9835 | 0.9778 |
| ncGTW | 0.9998 | 0.9998 |
| **Peak2** | | |
| Scores / Methods | MCC | SP |
| Before alignment | 0.1165 | 0.1937 |
| DBA | 0.7885 | 0.7957 |
| CPM | 0.3988 | 0.4801 |
| GTW | 0.9301 | 0.9379 |
| ncGTW | 0.9999 | 0.9998 |
| **Peak3** | | |
| Scores / Methods | MCC | SP |
| Before alignment | 0.0859 | 0.2002 |
| DBA | 0.7521 | 0.9999 |
| CPM | 0.3408 | 0.5831 |
| GTW | 0.6926 | 0.9407 |
| ncGTW | 0.7481 | 0.9998 |

**Table S2.** Peak scores for four methods of block structure.

| Peak1 | | |
|---|---|---|
| Scores / Methods | MCC | SP |
| Before alignment | 0.0686 | 0.2293 |
| DBA | 0.5475 | 0.9996 |
| CPM | 0.5931 | 0.9090 |
| GTW | 0.5312 | 0.8433 |
| ncGTW | 0.6750 | 0.9991 |
| **Peak2** | | |
| Scores / Methods | MCC | SP |
| Before alignment | 0.0695 | 0.2545 |
| DBA | 0.2809 | 0.7712 |
| CPM | 0.5041 | 0.8091 |
| GTW | 0.4759 | 0.8482 |
| ncGTW | 0.6673 | 0.9989 |
| **Peak3** | | |
| Scores / Methods | MCC | SP |
| Before alignment | 0.0542 | 0.2184 |
| DBA | 0.3108 | 0.5238 |
| CPM | 0.5980 | 0.9419 |
| GTW | 0.4707 | 0.8427 |
| ncGTW | 0.7435 | 0.9990 |

**Table S3.** Peak scores for four methods of non-informative structure.

### Supplementary Figure Legends

**Figure. S1.** Figures of DTW grids, DTW graphs, structural information diagram for GTW, and GTW graph. (a) A DTW grid for aligning $x_i$ to $x_j$. The purple path and the orange path correspond to two different warping functions. Each node in the grid is corresponds to a pair of points, one from $x_i$ and another from $x_j$. For example, node (3, 2) corresponds to the third point on $x_i$ ($x_{i3}$) and the second point on $x_j$ ($x_{j2}$). The weight of an edge is determined by its starting node. For example, the weight of the edge ((3, 2), (3, 3)) is given by the distance between $(x_{i3}, x_{j2})$. The distance between the purple path and the orange path is the area in dark blue (four triangles in this case). The corresponding warping function of the purple path is {(1, 1), (2, 2), (3, 2), (3, 3), (4, 4)}. (b) A DTW grid and the corresponding DTW graph, where the blue lines and dots form the original DTW grid, and the red lines and dots form the corresponding DTW graph. (c) An example of structural information between samples and the induced structure information between pairwise warping functions. Suppose there are five LC-MS samples and the run orders of which are exactly 1, 2, 3, 4, and 5. These samples are expected to have continuous changing profiles from smaller indices to larger indices. More importantly, the continuous changing give rise to the similarities between warping function. For example, the warping function for curve pair $(x_1, x_2)$ is similar to the warping function for $(x_2, x_3)$. In general, curve pairs $(x_i, x_{i+1})$ and $(x_{i+1}, x_{i+2})$ are considered as neighbors. Similarly, curve pairs $(x_i, x_j)$ and $(x_i, x_{j+1})$, along with curve pairs $(x_i, x_j)$ and $\left(x_{i+1}, x_j\right)$ are also neighbors. The diagram shows all warping function neighboring information. (d) Two small DTW graphs connected together. Green lines are the additional edges linking the corresponding vertices of the DTW graphs. (e) A GTW graph formed by two linked DTW graphs. Two warping functions ($\Phi_{i,j}$ and $\Phi_{k,l}$) are neighbors. Orange edges connect vertices to source and sink. Green edges link the corresponding vertices in two graphs. For clarity, we only show links between top three vertices. Other vertices are linked in the same way.

**Figure S2.** The illustration of stage 2 of ncGTW. (a) Example of inconsistency and nondiagonality calculation. From the DTW grid of the upper-left corner, to achieve consistency, the 2nd point on $x_i$ and the 3rd point on $x_j$ should be aligned to the same position on the virtual reference, because $(2, 3) \in \Phi_{i,j}$. It is clearly not the case by looking at the other two DTW grids. From $\Phi_{i,c}$ (lower-left DTW grid), we know that the 2nd point on $x_i$ is aligned to point (2, 3, 4) on the reference. From

$\Phi_{c,j}$ (upper-right DTW grid, the inverse of $\Phi_{j,c}$), we know that the 3rd point on $x_j$ is aligned to point (1, 2) on the reference. Thus, by definition the inconsistency from (2, 3) is $|4-2| + |2-1| = 3$. The total inconsistency is calculated along all nodes in $\Phi_{i,j}$. The nondiagonality of $\Phi_{i,c}$ is 6, since there are totally 6 corresponding vertical and horizontal paths in the DTW grid. The nondiagonality of $\Phi_{i,j}$ is 2. (b) The graph of stage 2 of ncGTW that solves the function. From stage 1, we have all pairwise warping functions. If from the warping function $\Phi_{i,j}$, we know that 2nd in $x_i$ is aligned to 3rd in $x_j$, this graph shows how to link the related vertices, where $x_c$ is the virtual reference.

**Figure S3.** Example of five-curve alignment by ncGTW with line structure. (a) Five continuously changing samples: the first one is drawn in "o", second "+", third "*", fourth "-", fifth "x". The shifts between the directly neighboring samples are all two points. (b) The induced neighborhood structure between warping functions.

**Figure S4.** Case study on line structure. (a) The first sample. (b) The five synthetic samples are drawn in the order of "o", "+", "*", "-", and "x". The shifts between neighboring samples are all one. All samples contain three peaks but the fourth sample, the third peak of which is missing (the other two are pointed by the arrows). (c) Alignment result of DBA. (d) CPM. (e) GTW. (f) ncGTW.

**Figure S5.** The way to connect two blocks dataset (5 samples in each block). One can see there are three types of pairs: within the 1st group, between 2 groups, and within the second group.

**Figure S6.** Case study on block structure. (a) The first sample. (b) The eighth sample, the third peak of which is missing. (c) The first block of the dataset. The samples are drawn in the order of "o", "+", "*", "-", and "x". The shifts between neighboring samples are all one. (d) The second block of the dataset. The five samples are drawn as the previous subfigure. In the following subfigures, only the eighth sample with marks (triangle) and its peaks pointed by arrows. (e) All ten samples. The shift between the two blocks is seven points. (f) Alignment result of DBA. (g) CPM. (h) GTW. (i) ncGTW.

**Figure S7.** Case study on non-informative structure. (a) A sample without peak 2 (marked as triangles in the following subfigures). (b) All ten simulated samples. (c) Alignment result of DBA. (d) CPM. (e) GTW. (f) ncGTW.

**Figure S8.** Case study on real LC-MS dataset of ten samples. (a) One exemplary sample with nine clear peaks marked with index. The second peak (pointed by an arrow) was wrongly aligned by most methods except GTW and ncGTW. (b) All ten samples. (c) Result of DBA. (d) CPM (e) GTW (f) ncGTW.

**Figure S9.** Case study on real LC-MS dataset of twenty samples from two batches. (a) The first sample from the first block, in which there are three peaks. (b) The sixth sample from the second block, in which there are also three peaks. (c) The first block. For the third peak group, only two samples have the peak (pointed by the black arrows). (d) The second block. Only three samples have the peak in the third peak group (pointed by the gray arrows). (e) All twenty samples. (f) Result of DBA. (g) CPM (h) GTW (i) ncGTW.

**Figure S10.** Case study on real LC-MS dataset of ten samples but without the structure information. (a) One exemplary sample with nine clear peaks marked with index. The second peak (pointed by an arrow) was wrongly aligned by most methods except GTW and ncGTW. (b) All ten samples. (c) Result of DBA. (d) CPM (e) GTW (f) ncGTW.

**Figure S11.** Diagram of two-layer ncGTW

**Figure S12.** A XCMS aligned feature from the Rotterdam dataset as an example of false positives of the misalignment detection algorithm. One can see that all the peaks are aligned well. That is, unlike **Fig. 5b** in the main article, no peaks are spread consecutively across RT, Thus, this detected feature is considered as a false positive. The sample indexes in the red box are 34, 36, and 40, and the p-value is 0.034. Apparently, there are more than three peaks in the red box, but only three of them are detected. Moreover, the sample indexes in the blue box are 1, 4, 9, 21, 23, and 24, and the p-value is 0.047. Again, there are many peaks that are not detected in the blue box. Both of the p-values are lower than the threshold and the index sets are disjoint, so this feature is reported as misaligned.

#### References

Arnold, B.C., Balakrishnan, N. and Nagaraja, H.N. A first course in order statistics. Siam; 1992.

Connor, R.J. The Sampling Distribution of the Range from Discrete Uniform Finite Populations and a Range Test for Homogeneity. *Journal of the American Statistical Association* 1969;64(328):1443-1458.

Efron, B. and Morris, C. Stein's paradox in statistics. *Scientific American* 1977;236(5):119-127.

Jiang, W., *et al.* Comparisons of five algorithms for chromatogram alignment. *Chromatographia* 2013;76(17-18):1067-1078.

Korte, B., *et al.* Combinatorial optimization. Springer; 2012.

Listgarten, J., *et al.* Multiple alignment of continuous time series. In, *Advances in neural information processing systems*. 2005. p. 817-824.

Petitjean, F., Ketterlin, A. and Gançarski, P. A global averaging method for dynamic time warping, with applications to clustering. *Pattern Recognition* 2011;44(3):678-693.

Wang, Y., *et al.* Graphical time warping for joint alignment of multiple curves. In, *Advances in Neural Information Processing Systems*. 2016. p. 3648-3656.

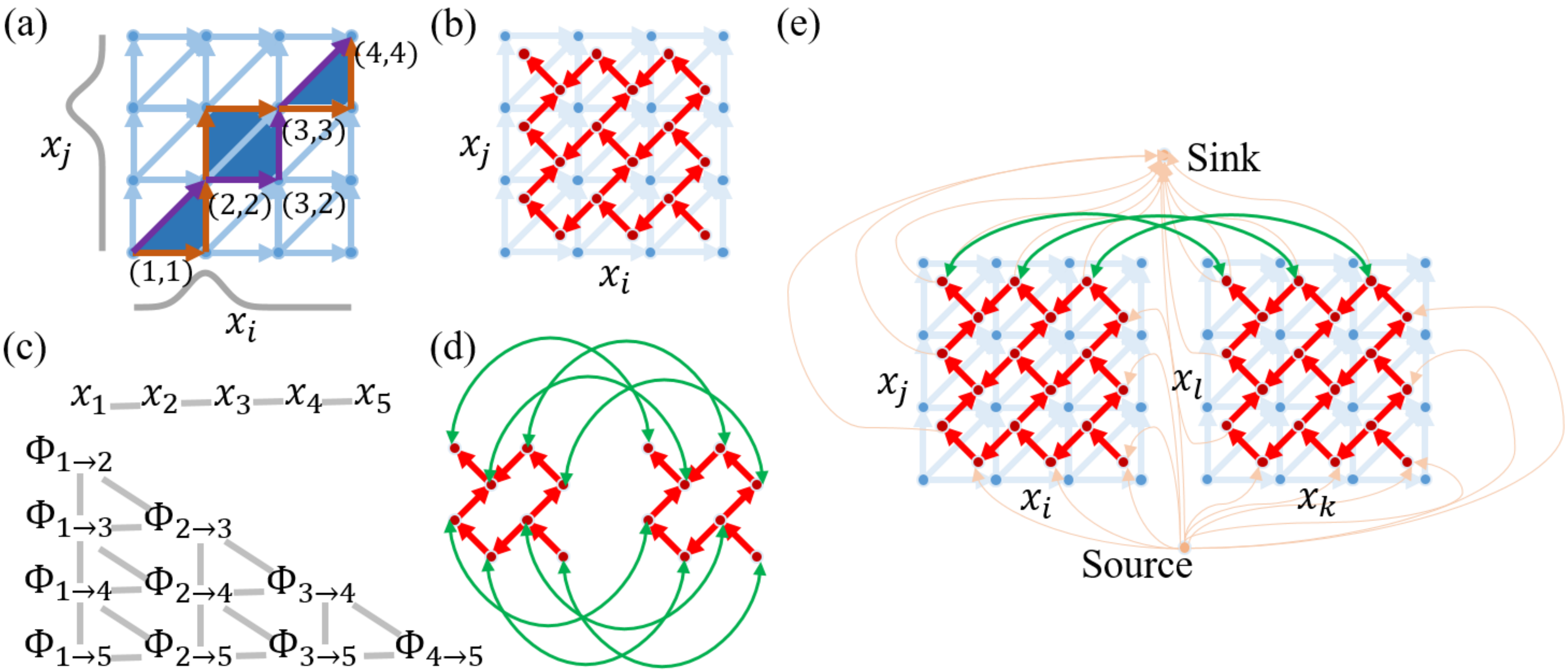

(a)
(4,4)
(3,3)
(2,2)
(3,2)
(1,1)
$x_j$
$x_i$
(b)
$x_j$
$x_i$
(e)
Sink
$x_j$
$x_l$
$x_i$
$x_k$
Source
(c)
$x_1$ $x_2$ $x_3$ $x_4$ $x_5$
$\Phi_{1\to2}$
$\Phi_{1\to3}$ $\Phi_{2\to3}$
$\Phi_{1\to4}$ $\Phi_{2\to4}$ $\Phi_{3\to4}$
$\Phi_{1\to5}$ $\Phi_{2\to5}$ $\Phi_{3\to5}$ $\Phi_{4\to5}$
(d)

Fig. S1

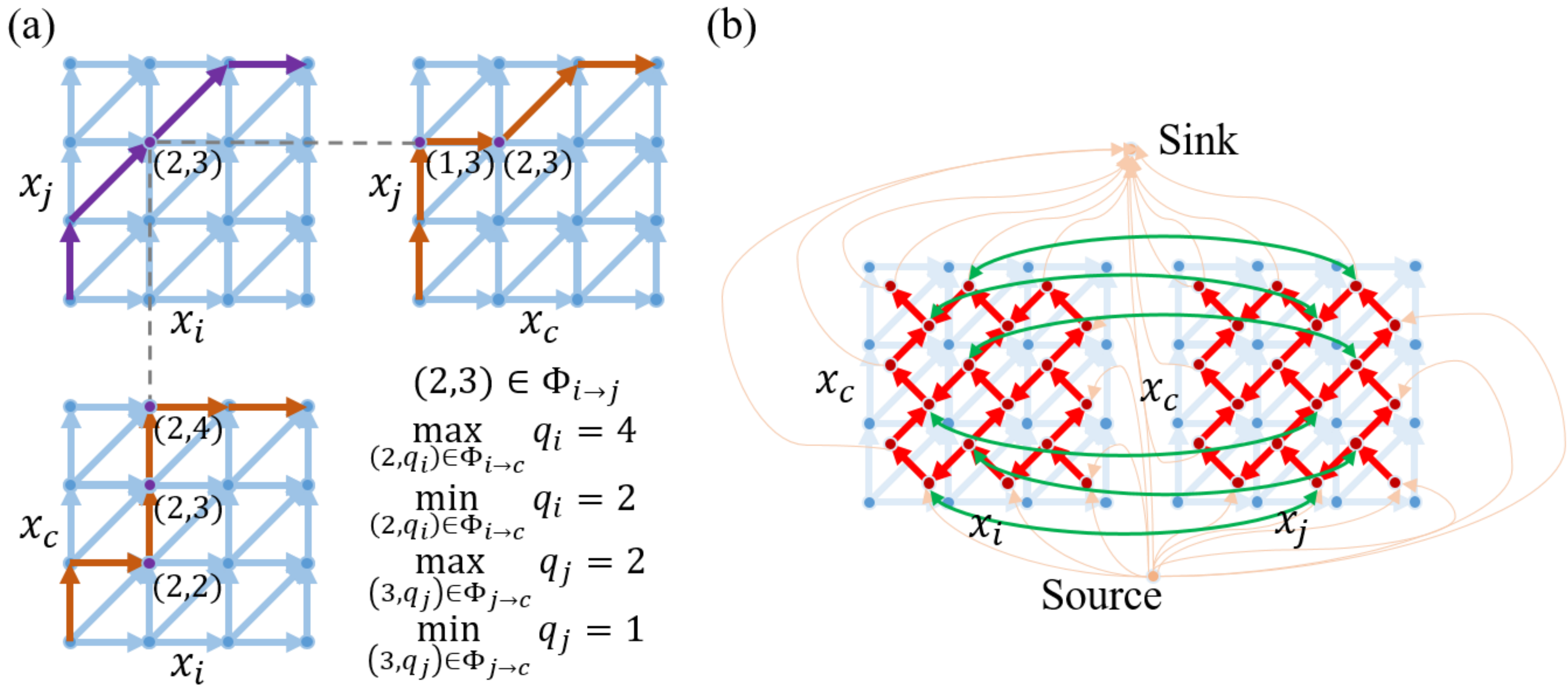
(a)
(b)
(2,3)
(1,3)
(2,3)
(2,4)
(2,3)
(2,2)
$x_j$
$x_i$
$x_c$
$(2,3) \in \Phi_{i \to j}$
$\max_{(2,q_i) \in \Phi_{i \to c}} q_i = 4$
$\min_{(2,q_i) \in \Phi_{i \to c}} q_i = 2$
$\max_{(3,q_j) \in \Phi_{j \to c}} q_j = 2$
$\min_{(3,q_j) \in \Phi_{j \to c}} q_j = 1$
Sink
Source

Fig. S2

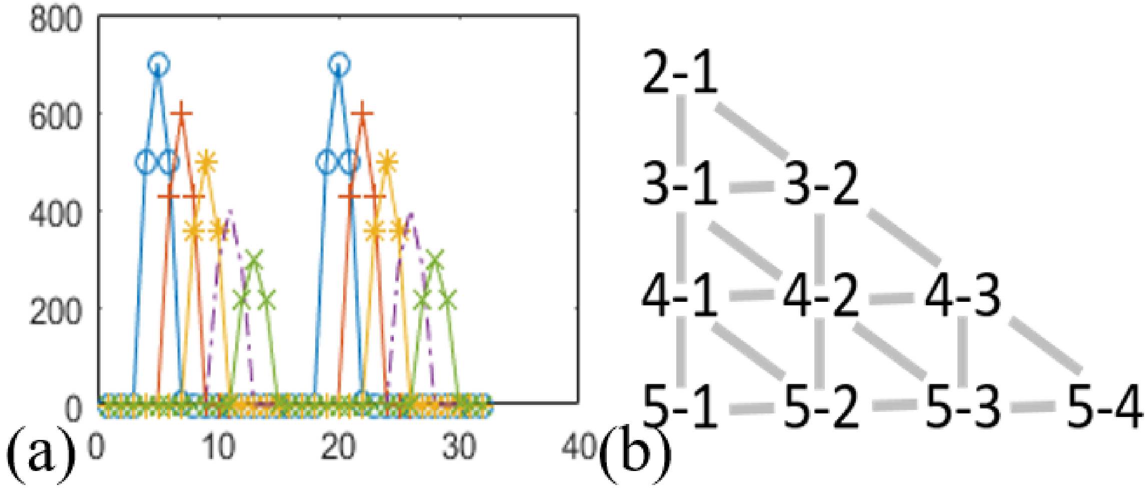

Fig. S3

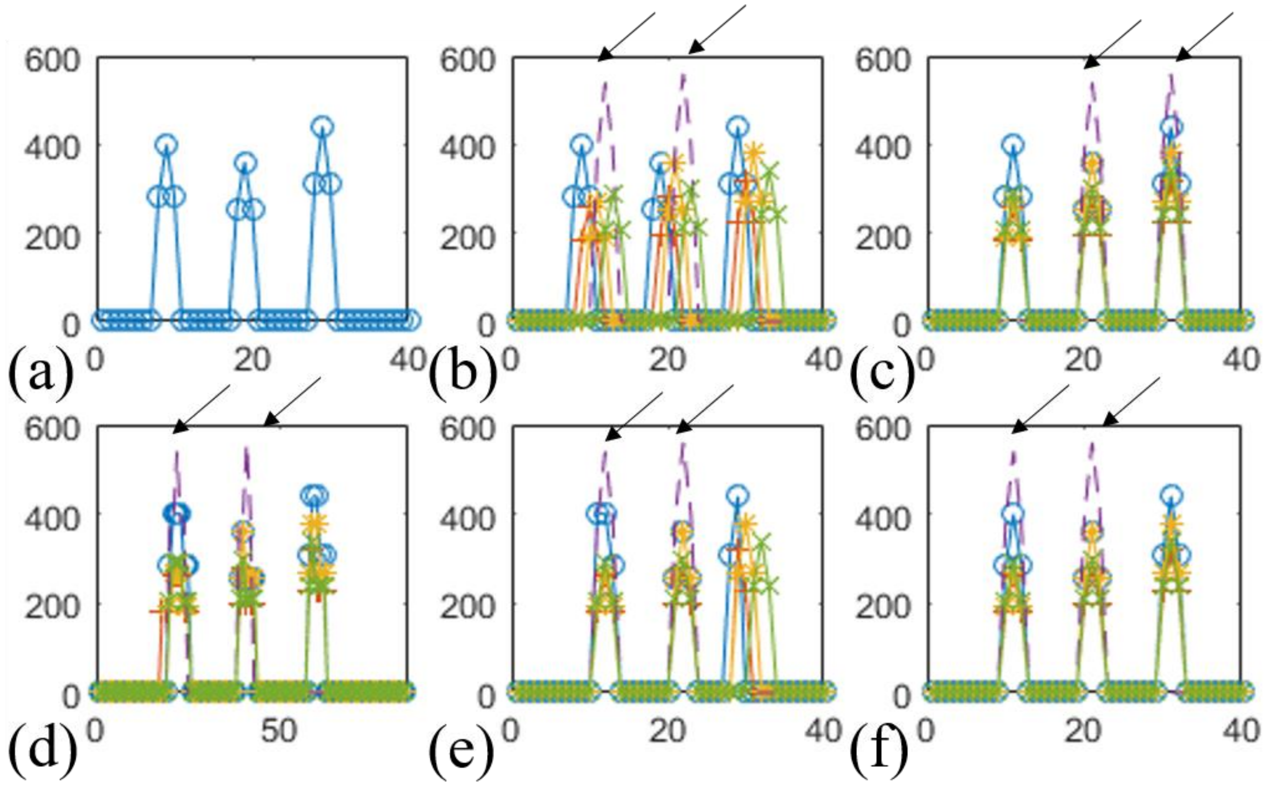

Fig. S4

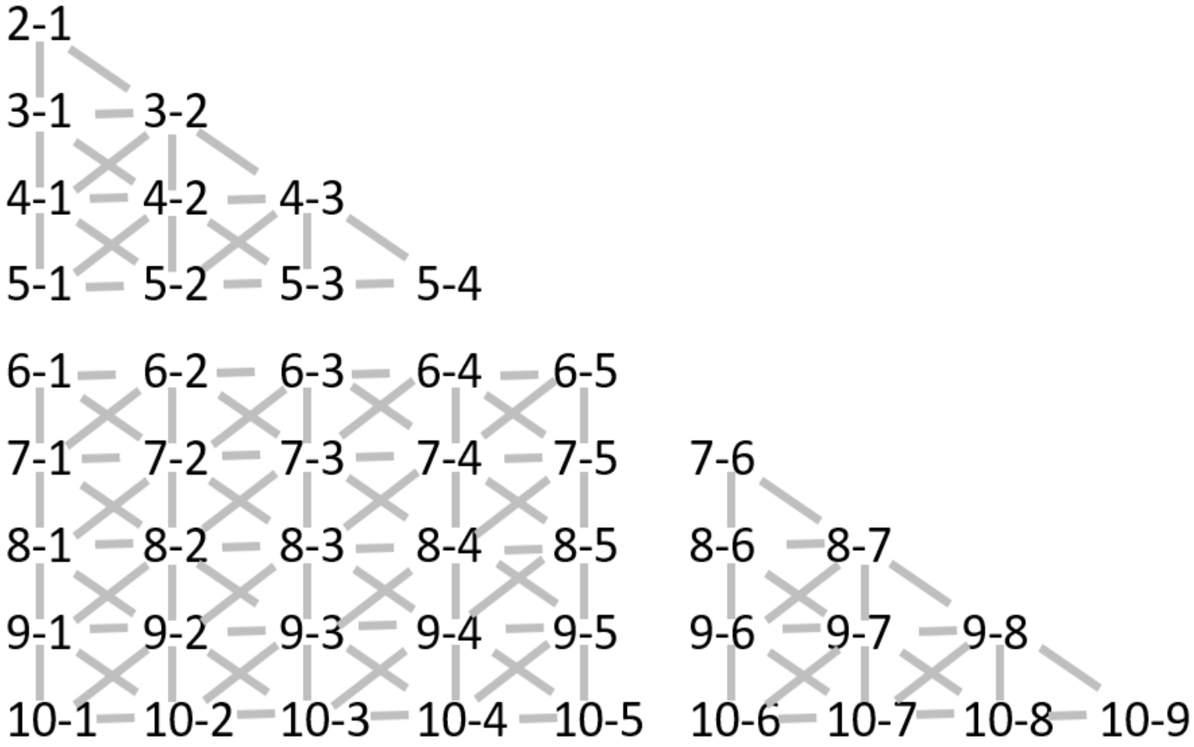
2-1
3-1 3-2
4-1 4-2 4-3
5-1 5-2 5-3 5-4
6-1 6-2 6-3 6-4 6-5
7-1 7-2 7-3 7-4 7-5 7-6
8-1 8-2 8-3 8-4 8-5 8-6 8-7
9-1 9-2 9-3 9-4 9-5 9-6 9-7 9-8
10-1 10-2 10-3 10-4 10-5 10-6 10-7 10-8 10-9

Fig. S5

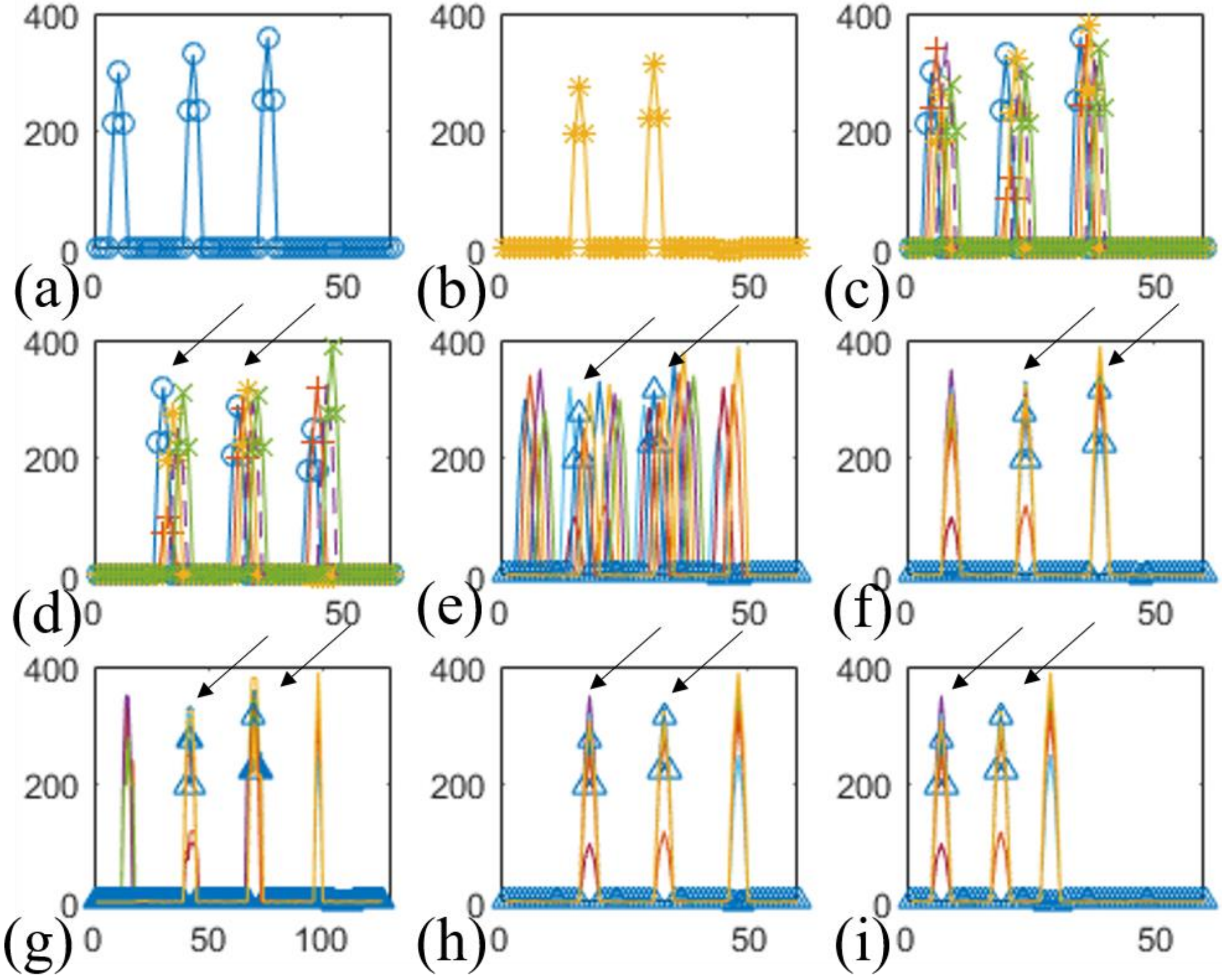

Fig. S6

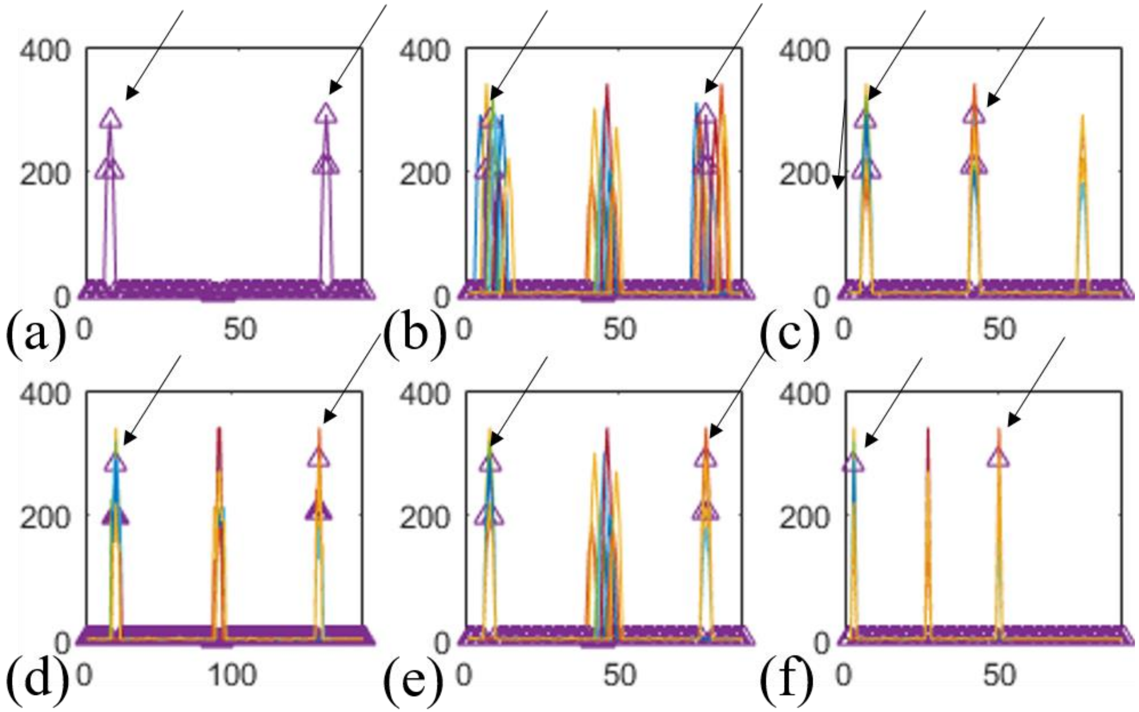

Fig. S7

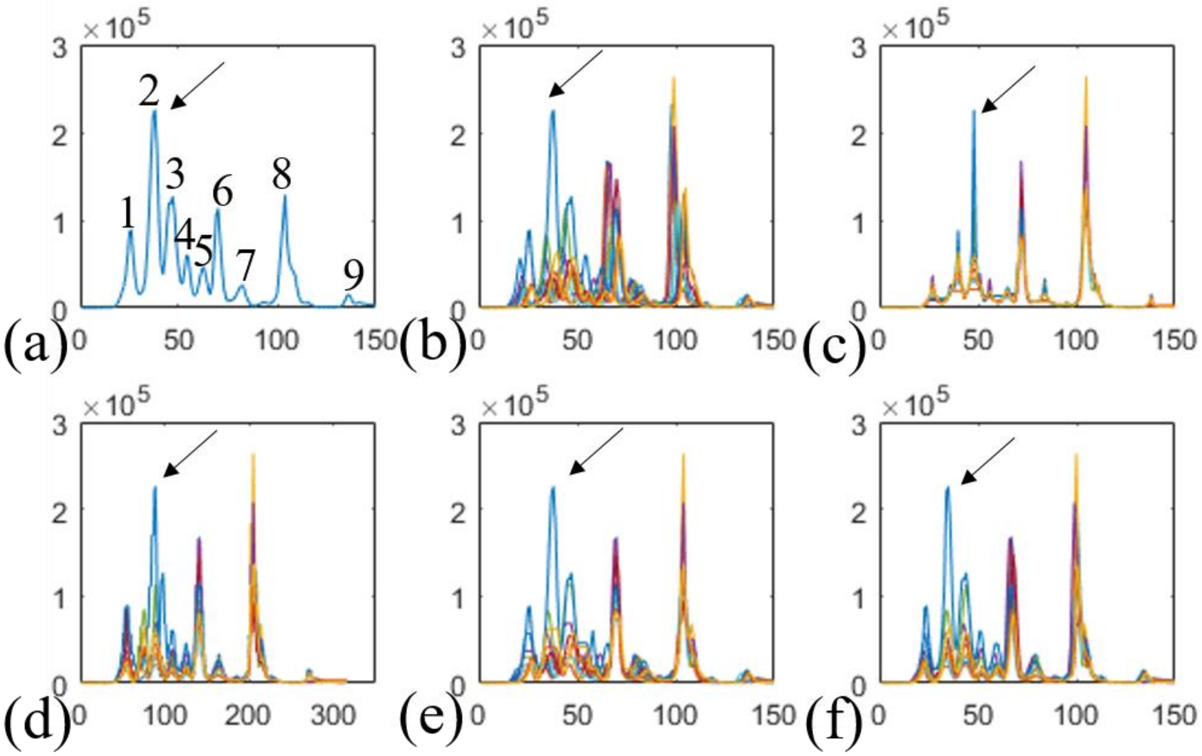

Fig. S8

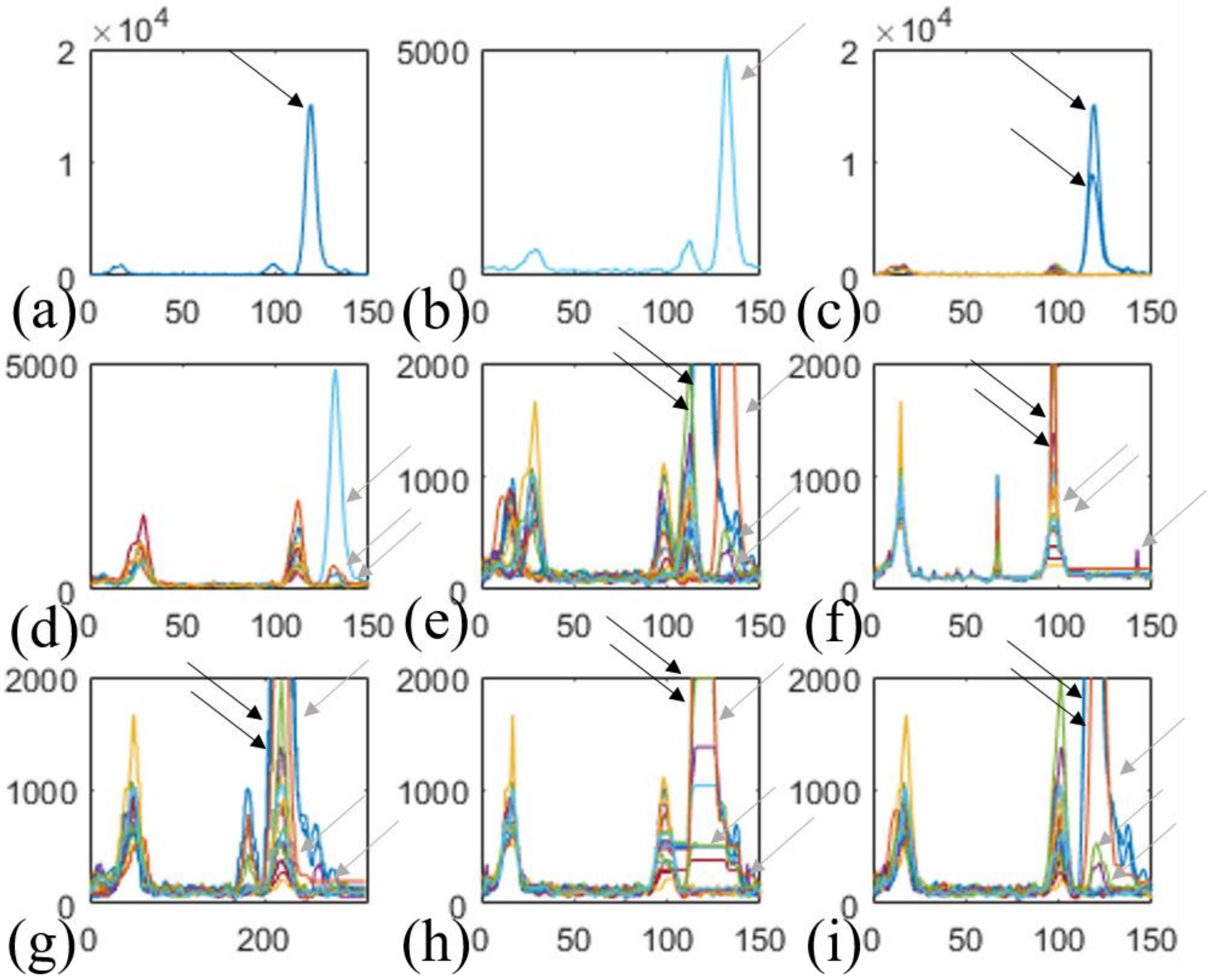
×10^4
2
1
0
(a)
0 50 100 150
5000
0
(b)
0 50 100 150
(c)
(d)
2000
1000
(e)
(f)
(g)
0 200
(h)
(i)

Fig. S9

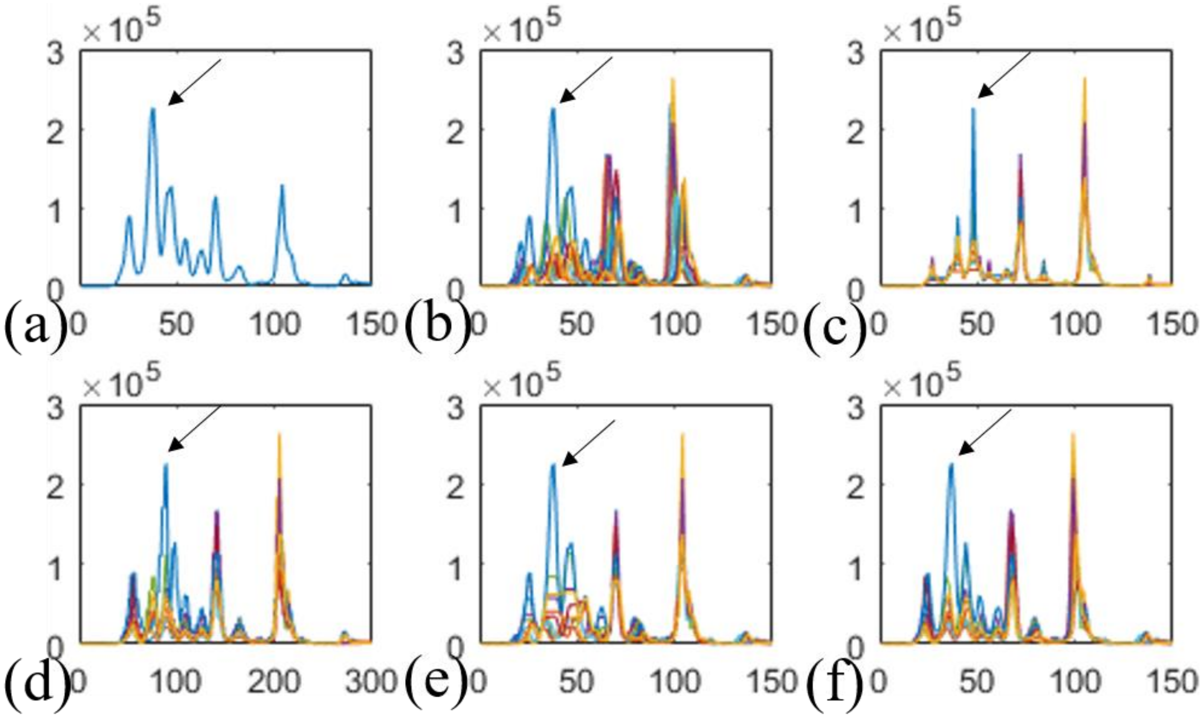

×10^5
3
2
1
0
(a)
0 50 100 150
(b)
0 50 100 150
(c)
0 50 100 150
(d)
0 100 200 300
(e)
0 50 100 150
(f)
0 50 100 150

Fig. S10

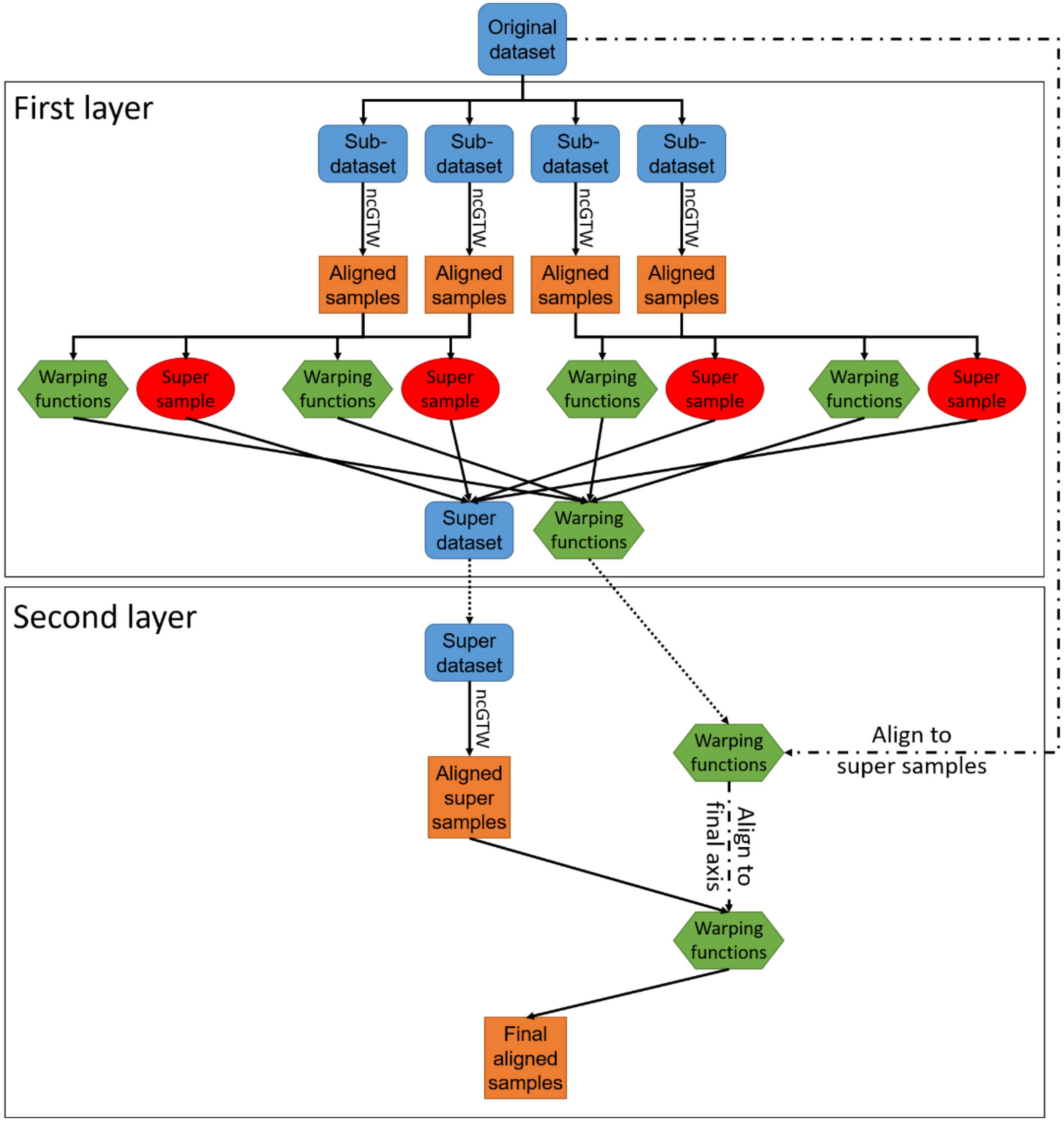
Original dataset
First layer
Sub-dataset
Sub-dataset
Sub-dataset
Sub-dataset
ncGTW
ncGTW
ncGTW
ncGTW
Aligned samples
Aligned samples
Aligned samples
Aligned samples
Warping functions
Super sample
Warping functions
Super sample
Warping functions
Super sample
Warping functions
Super sample
Super dataset
Warping functions
Second layer
Super dataset
ncGTW
Aligned super samples
Warping functions
Align to super samples
Align to final axis
Warping functions
Final aligned samples

Fig. S11

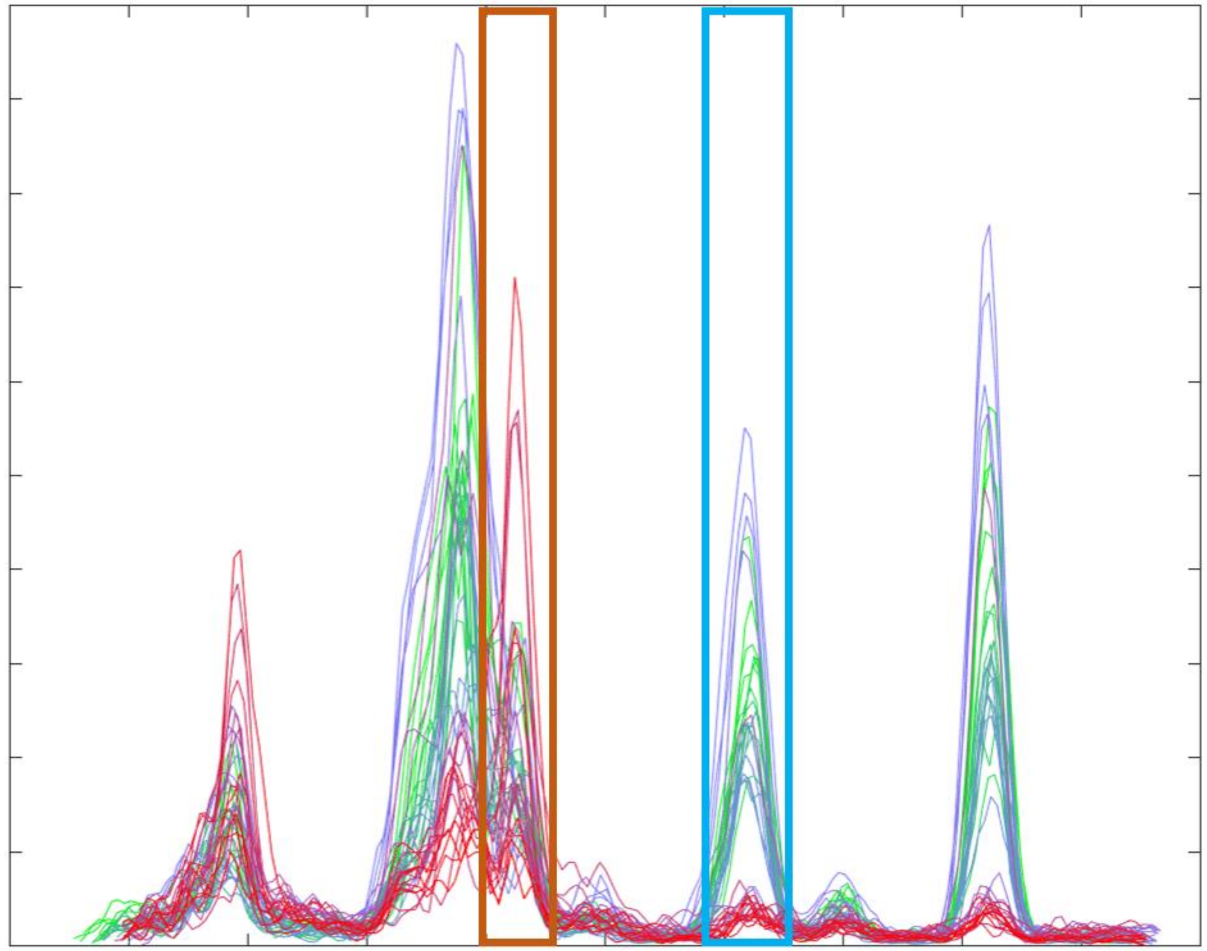

Fig. S12
